# Transcriptomic learning for digital pathology

**DOI:** 10.1101/760173

**Authors:** Benoît Schmauch, Alberto Romagnoni, Elodie Pronier, Charlie Saillard, Pascale Maillé, Julien Calderaro, Meriem Sefta, Sylvain Toldo, Mikhail Zaslavskiy, Thomas Clozel, Matahi Moarii, Pierre Courtiol, Gilles Wainrib

## Abstract

Deep learning methods for digital pathology analysis have proved an effective way to address multiple clinical questions, from diagnosis to prognosis and even to prediction of treatment outcomes. They have also recently been used to predict gene mutations from pathology images, but no comprehensive evaluation of their potential for extracting molecular features from histology slides has yet been performed. We propose a novel approach based on the integration of multiple data modes, and show that our deep learning model, HE2RNA, can be trained to systematically predict RNA-Seq profiles from whole-slide images alone, without the need for expert annotation. HE2RNA is interpretable by design, opening up new opportunities for virtual staining. In fact, it provides virtual spatialization of gene expression, as validated by double-staining on an independent dataset. Moreover, the transcriptomic representation learned by HE2RNA can be transferred to improve predictive performance for other tasks, particularly for small datasets. As an example of a task with direct clinical impact, we studied the prediction of microsatellite instability from hematoxylin & eosin stained images and our results show that better performance can be achieved in this setting.

---

Histological analyses of tumor biopsy sections have been important tools in oncology for more than a century, providing a high-resolution map of the tumor that helps pathologists to determine both diagnosis and grade^1,2^. The development of new, more powerful technologies, and the curation of larger datasets, have made it possible to train increasingly sophisticated algorithms, which can process and learn from very high-definition whole-slide digital images (WSI). Deep convolutional neural networks (CNNs) have recently emerged as an important image analysis tool, accelerating the work of pathologists. They have shattered performance benchmarks in many challenging medical applications, including mitosis detection^3,4^, the quantification of tumor immune infiltration^5^, cancer subtypes classification^6,7^ and grading^8,9^. Ultimately, they enhance the practices of pathologists, improving the prediction of patient survival outcomes and response to treatment^10,11^ and presenting exciting opportunities in clinical and biomedical fields^12,13^.

However, while it is becoming clear that the application of deep learning models to tissue-based pathology can be very useful, few attempts have been made to connect specific molecular signatures directly to morphological patterns within cancer subtypes. Several recent studies have shown how models of this class can connect histological images to tumor-specific mutations or tumor mutational burden in lung^10^, prostate^14^, brain cancers^15^ and melanoma^16,17,18^. Massive changes in gene expression are known to occur in many human cancers secondary to activating/silencing mutations or epigenomic modifications, and the comprehensive characterization of disease-related gene networks/signatures can help to clarify potential disease mechanisms and prioritize targets for novel therapeutic approaches^19,20^. Various next-generation sequencing techniques have been developed for the reconstruction of gene information in carcinogenesis^21,22^, together with specific bioinformatic tools for their analysis^23,24,25^. However, despite the continual decrease in the cost of such next-generation sequencing^26^, these technologies are not routinely used by all medical centers. Moreover, they are associated with several challenges, such as the need for sufficiently high DNA quality and quantity, and they are time-consuming and difficult to incorporate into routine clinical practice. An ability to predict gene expression from histology slides would therefore greatly facilitate patient diagnosis and prediction of response to treatment and survival outcome^27^.

We have begun to fill this gap, with the development of HE2RNA, a new deep learning algorithm, specifically customized for the direct prediction of gene expression from WSI (Fig. 1). For the training and testing of our model, we collected WSIs for which the corresponding RNA-Seq data were available, from the public dataset The Cancer Genome Atlas (TCGA). We then investigated how HE2RNA learned to recognize important histological patterns within the slides, by using our model to generate heatmaps corresponding to the most predictive tiles used to predict a gene’s expression. Such approaches could also be used to detect histological subtypes^10^, genetic mutations^28^ or to map the infiltration of tumors by tumor-infiltrating lymphocytes (TILs)^29^ or other immune cells (e.g. macrophages, NK cells) based on cell-specific gene signatures. Such models have also been used to predict molecular profiling from pathology slides, particularly for the prediction of the hormonal status of breast cancer cases^30^.

**Fig. 1:**
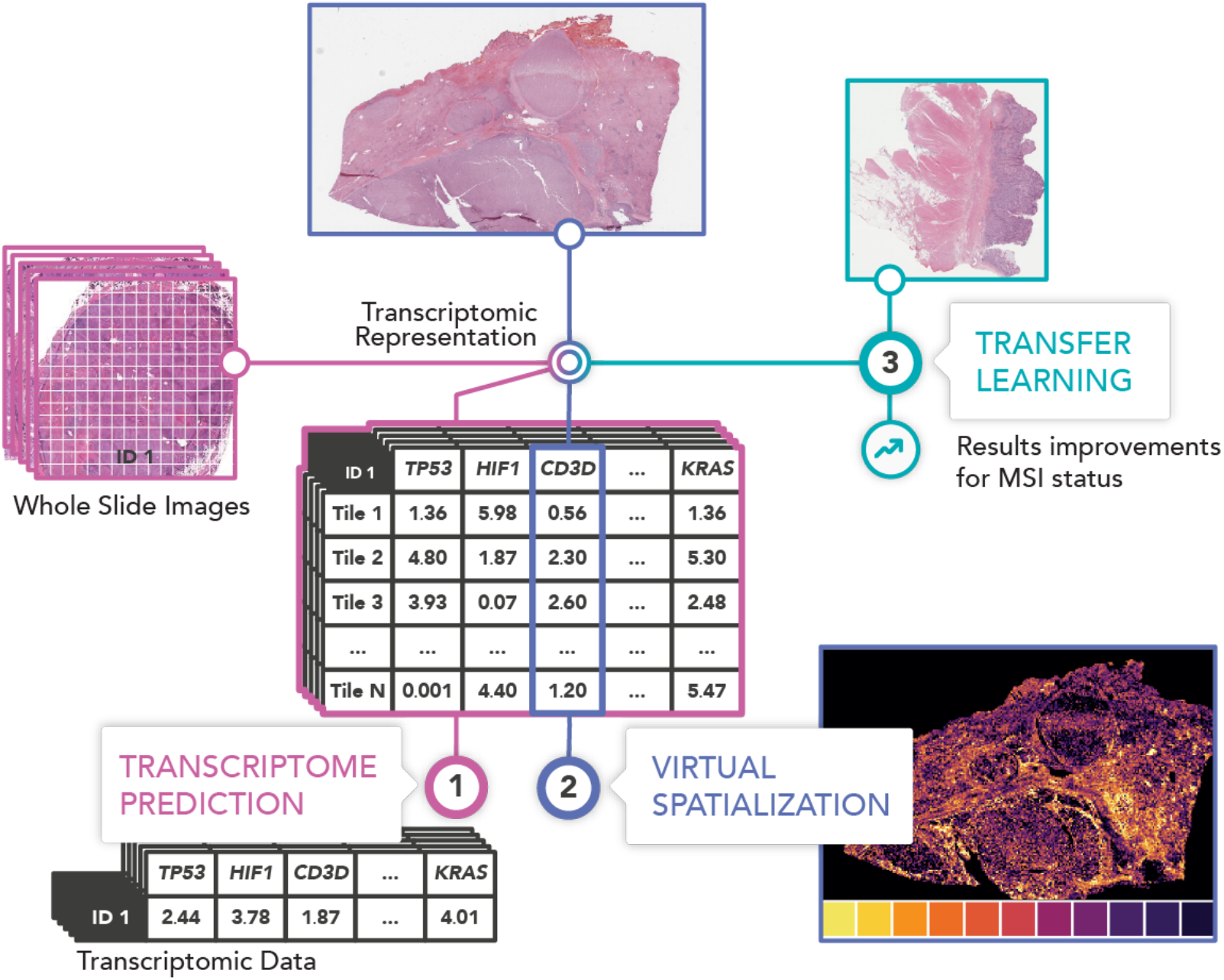
Graphical abstract: Transcriptomic learning for digital pathology. Hematoxylin & eosin (H&E)-stained histology slides and RNA-Seq data (FPKM-UQ values) for 28 different cancer types and 8,725 patients were collected from The Cancer Genome Atlas (TCGA) and used to train the neural network HE2RNA to predict transcriptomic profile from the corresponding high-definition whole-slide images (WSI). During this task, the neural network learned an internal representation encoding both information from tiled images and gene expression levels. This *transcriptomic representation* can be used for: **1. Transcriptome prediction** from images without associated RNA sequencing. **2. The virtual spatialization of transcriptomic data.** For each predicted coding or non-coding gene a score is calculated for each tile on the corresponding WSI. These predictive scores can be used to generate heatmaps for each gene for which expression is significantly predicted. **3. Improving predictive performances for different tasks, in a transfer learning framework,** as shown here for a realistic setup, for microsatellite instability (MSI) status prediction from non-annotated WSIs.

Finally, the information linking the WSI to the RNA-Seq data, extracted during transcriptomic learning on a large dataset, such as TCGA, and encoded in the internal representation of the model, which we refer to as *transcriptomic representation*, could be transferred to improve the performance of other predictive tasks, such as the prediction of microsatellite instability (MSI) status. We validated our hypothesis in a realistic setup, in which only a small dataset was available for the learning phase of the MSI status, built directly from histological images obtained from patients with colorectal adenocarcinomas (COAD), although a *transcriptomic representation* was available from a larger dataset of COAD patients. This aspect is particularly important, as it has recently been demonstrated that MSI-H (MSI-High) status is a good predictive biomarker for potential responders to immunotherapy, resulting in a better overall prognosis relative to that of patients with microsatellite stable disease (MSS), in gastric adenocarcinoma (STAD) and colorectal cancer^31^. However, not all patients are screened for MSI status outside of high-volume tertiary care centers. The possibility of inferring MSI status directly from biopsies would therefore potentially improve the diagnosis and medical care of this subgroup of patients.

## Results

### A deep learning model for the prediction of gene expression

Gene expression is highly variable and influenced by cell-type, proliferation and differentiation status^32,33^. However, the possibility of quantifying the expression levels of specific genes based on a visual observation of hematoxylin & and eosin (H&E)-stained WSI from tissue biopsies has never been investigated in detail. We aimed to tackle this problem, for multiple cancers, by developing HE2RNA, a deep learning model based on a multi-task weakly-supervised multiple-instance learning approach^34^ (architecture explained in the Methods section). We used data from 8,725 patients with matched WSIs and RNA-Seq profiles, corresponding to 28 different cancer types, from the TCGA network (see Methods for details about the dataset). The model was trained to use WSIs as inputs to predict FPKM-UQs (the outputs). We performed a 5-fold cross-validation strategy. The slides were thus randomly assigned to five different sets, and each set was used in turn as the validation set, the other four sets being used for training in the run concerned. For each cancer type, the patients were split 80%-20% between the training/validation sets in each run and the final results are expressed as a mean for all five runs (see the Methods section for more details).

For RNA-Seq data-preprocessing, we selected the 30,839 (coding or non-coding) genes with positive median FPKM-UQ values and considered their expression levels on a logarithmic scale, using an approach similar to that used in existing bioinformatic tools^35^ for differential gene expression analysis (see Methods for more detail). WSIs are high-definition digital images (up to 100,000 × 100,000 pixels). We therefore preprocessed the slides, to facilitate the development of our model, by partitioning them into “tiles”, corresponding to small squares of 112 × 112 μm (224 × 224 pixels). The tiles were then aggregated into clusters, called super-tiles. The number and size of the super-tiles were optimized, for each specific task, at the preprocessing level. In particular, for the transcriptome prediction task, 100 super-tiles were created for each WSI. A multilayer perceptron was then applied to the ensemble of super-tiles to generate a predicted value per gene and per super-tile. For comparison of the model predictions with the real RNA-Seq data, the predictions per-super-tile were aggregated by calculating a weighted mean to give a final prediction per-WSI (see details in Methods and Supplementary Fig. S1).

Finally, we evaluated the results for each gene, by calculating Pearson’s coefficient *R* for the correlation over samples between the model predictions and the real data. This correlation was assessed for each preselected gene, separately for each different type of cancer. We considered a prediction for a given gene to be significantly different from the random baseline value if the *p*-value associated with its coefficient *R* was below 0.05, after correction to account for the testing of multiple hypotheses. We considered Holm-Šidák (HS) correction and Benjamini-Hochberg (BH) correction, with significance level *α*=0.05 in each case. For the total set of 30,839 selected genes, an average of 3,627 genes (including 2,797 protein-coding) per cancer type were predicted with a statistically significant correlation under HS correction, whereas an average of 12,853 genes (including 8,450 protein-coding genes) were predicted with a statistically significant correlation following BH adjustment. The results shown in Fig. 2 and discussed in the text were obtained with the more conservative HS method. The results obtained with BH adjustment are shown in Supplementary Fig. S2.

**Fig. 2:**
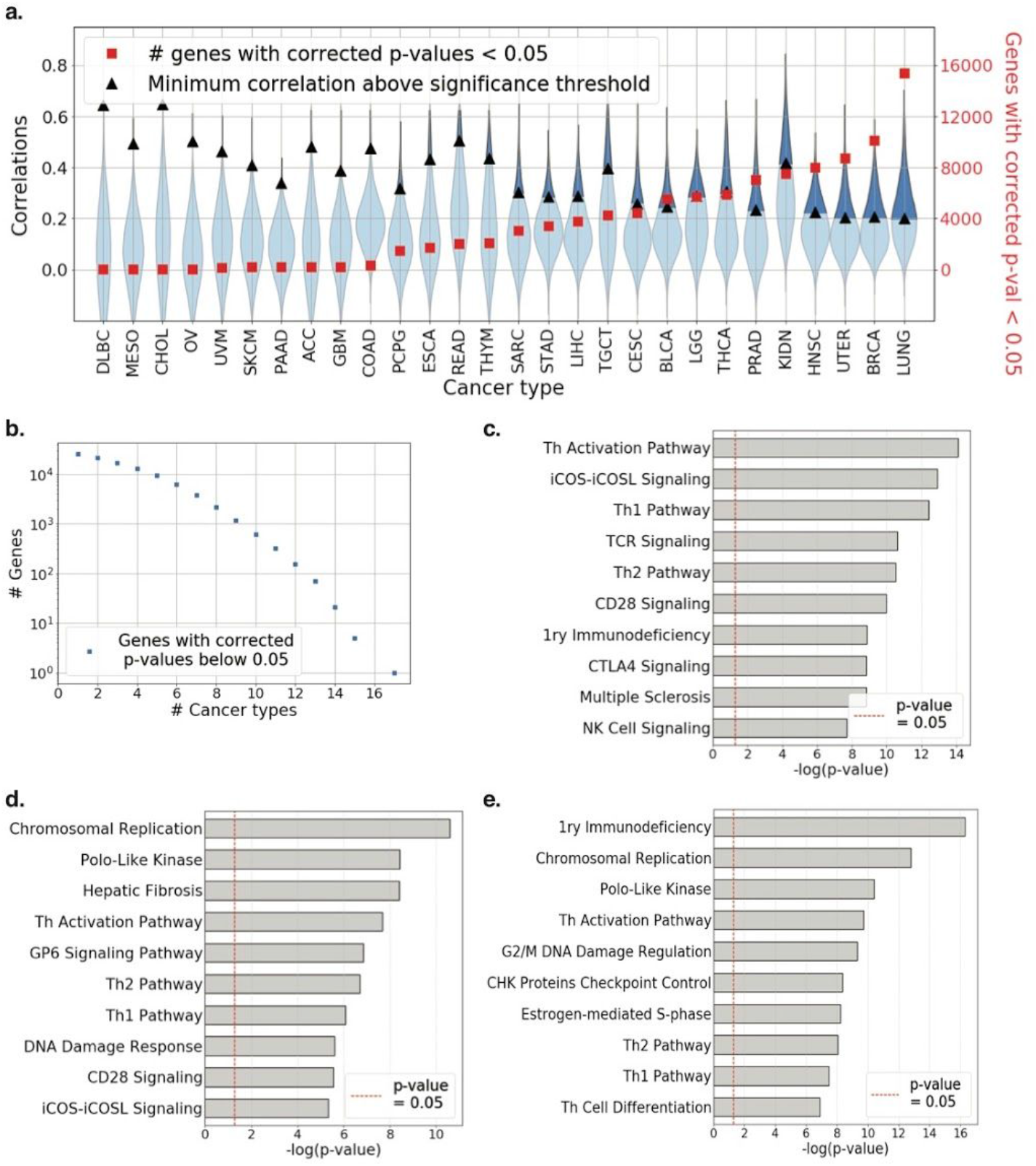
Gene expression prediction results. **a.** Distribution of Pearson correlation coefficients *R* (left axis, blue violin plots) and the number of coding and non-coding genes (right axis, red squares) with Holm-Šidák corrected *p*-values <0.05, for 28 cancer types from the TCGA. Black triangles indicate the minimum correlation coefficient required for significance in any given dataset. **b.** Number of coding and non-coding genes significantly well-predicted for a given number of cancer types, as a function of the number of cancers. **c.** Computational pathway analysis with Ingenuity Pathway Analysis (IPA) software of the 156 best-predicted Pan TCGA genes, showing an enrichment in genes associated with immunity and tumor immune infiltration/activity. TCR: T cell receptor, NK: natural killer. **d.** IPA-based analysis of the more accurately predicted protein-coding genes in the LIHC dataset, showing an enrichment in genes associated with cell cycle and DNA damage response. LIHC: liver hepatocellular carcinoma. **e.** IPA-based analysis of the more accurately predicted protein-coding genes in the BRCA dataset, showing an enrichment in genes associated with cell cycle and DNA damage response. Th Cell differentiation: Th1 and Th2 cell differentiation. BRCA: breast cancer. Th activation pathway: Th1 and Th2 activation pathway, 1ry: Primary.

The number of significantly well-predicted genes varied considerably between cancer types, mostly due to the size of the dataset considered (Fig. 2a): the smaller the number of samples in the dataset, the higher the correlation coefficient *R* required for statistical significance. For example, only seven genes were accurately predicted for the 44 available cases of Lymphoid Neoplasm Diffuse Large B-cell Lymphoma (DLBC), (*R* > *R*_*sign*_ = 0.64), whereas 15,391 genes were correctly predicted for the 1,046 cases of lung carcinoma (indicated as LUNG, including 535 WSIs for lung adenocarcinoma - LUAD - and 511 slides for lung squamous cell carcinoma - LUSC), (*R* > *R_sign_* = 0.20). The correlation coefficients obtained for the well-predicted genes were often well above the threshold value for significance threshold, particularly for the largest datasets (Fig. 2a).

We compared the list of genes well-predicted in each cancer, to analyze the consistency of the predictions. None of the genes were well-predicted in all 28 available cancer types (Fig. 2b), but few genes were consistently above the significance threshold when considering smaller subsets of cancer. In particular, *C1QB* expression was strikingly well-predicted in 17/28 different cancer datasets (*R* = 0.39 ± 0.15). Similarly, another four genes, *NKG7*, *ARHGAP9*, *C1QA* and *CD53* were accurately predicted in 15/28 datasets (*R* ranging from 0.38 to 0.46 for the various cancer types). C1Q (A and B) are proteins of the complement known to be involved in T-cell activation following antigen presentation by antigen presenting cells (APC)^36^, whereas CD53^37^ and NKG7^38^ are known to be expressed by T and NK cells, respectively.

Longer lists of genes were consistently well-predicted by HE2RNA in smaller subsets of cancer types, and we used Ingenuity Pathway Analysis (IPA) software to identify the corresponding biological networks. For example, we performed a functional annotation of the 156 genes with prediction performances above the significance threshold in 12/28 different cancer types (Fig. 2c).

This analysis revealed an enrichment in genes involved in the immune system and T-cell regulation/activation in the various cancer types, as already suggested by the genes mentioned above. Indeed, the most significant functional network was the Th1 and Th2 activation pathway (p-value = 7.94 × 10^−15^), and other networks with similar levels of significance included iCOS-iCOSL signaling in T helper cells, T-cell receptor signaling and CD28 signaling in T helper cells. Gene expression profiles differ considerably between types of cancer. We therefore performed a similar analysis on two specific examples, liver hepatocellular carcinomas (LIHC) and invasive breast carcinoma (BRCA). In LIHC the genes for which expression was most accurately predicted were associated with mitosis and cell-cycle control (Cell cycle control of chromosomal Replication, Mitotic Roles of Polo-Like Kinase), known hallmarks of cancer (Fig. 2d). Liver fibrosis, a known risk factor for the development of LIHC^39^, was also among the most significantly well-predicted networks (Hepatic Fibrosis/Hepatic Stellate Cell Activation network). Similarly, we found that the model performed well on BRCA samples (Fig. 2e), for the prediction of expression levels for genes involved in cell-cycle regulation (Cell Cycle: G2/M DNA Damage Checkpoint Regulation, Cell cycle control of chromosomal Replication, Mitotic Roles of Polo-Like Kinase), but also for the prediction of expression levels for *CHEK2* (a gene known to be mutated in breast cancer^40^ and involved in its progression^41^) and *Cyclin E* (known to be overexpressed in breast cancer^42^). These results demonstrate that HE2RNA, despite training on a diverse range of cancer types, was able not only to predict expression levels for genes involved in immune regulation, but also to detect pathways deregulated in specific types of cancer.

Finally, we investigated whether known gene signatures dysregulated in a majority of cancer types and implicated in carcinogenesis could be accurately predicted by HE2RNA. We focused on the hallmarks of cancer corresponding to the six biological capabilities acquired during the multistep development of human tumors^43,44^: sustaining proliferative signaling, resisting cell death, evading growth suppressors, enabling replicative immortality, inducing angiogenesis, and activating invasion and metastasis. Based on these hallmarks, we defined a subset of genes known to be involved in the changes required for a normal cell to become cancerous: increased angiogenesis, increased hypoxia, deregulation of the DNA repair system, increased cell cycle activity, immune response mediated by B cells and adaptive immune response mediated by T cells. We combined several gene signatures from Gene Set Enrichment Analysis (GSEA) software for each of these biological networks, to obtain six gene signatures deregulated in several types of cancers (see Methods and full lists in Table S4).

For each of these pathways and for each cancer type, we compared the predictions obtained for the genes in the signatures with those obtained for 10,000 different random lists of genes of the same length. For each pathway length, we collected the corresponding distributions of gene prediction correlations *P* (*R*). We then compared (Fig. 3a) the mean correlation coefficient for the pathway *R*_*p*_ to the average of the mean correlation coefficients for the random lists *R*_*o*_. The model provided better predictions for all the pathways considered, with the vast majority of points lying above the identity line (Fig. 3a). Moreover, for each cancer type, we calculated the p-value required to obtain the signature correlation value *R*_*p*_ in the relative random distribution *P* (*R*). We found that, in 50% of cancer types for angiogenesis, and 54% for hypoxia, DNA repair and cell-cycle pathways, signatures were significantly better predicted by HE2RNA than random lists of genes (cumulating all levels of significance), with these proportions reaching 75% and 86% for B cell-mediated immunity and T cell-mediated immunity, respectively (Fig. 3a). We obtained similar results when comparing the proportion of well-predicted genes in cancer-specific pathways to the proportion of well-predicted genes in random sets of genes. In this case, the HE2RNA predictions were significantly better than those for a random set of genes in 36% (angiogenesis), 29% (hypoxia), 25% (DNA repair), 39% (cell-cycle), 36% (B cells-mediated immunity) and 50% (T cells-mediated immunity) of cancer types (Fig. 3b).

**Fig. 3:**
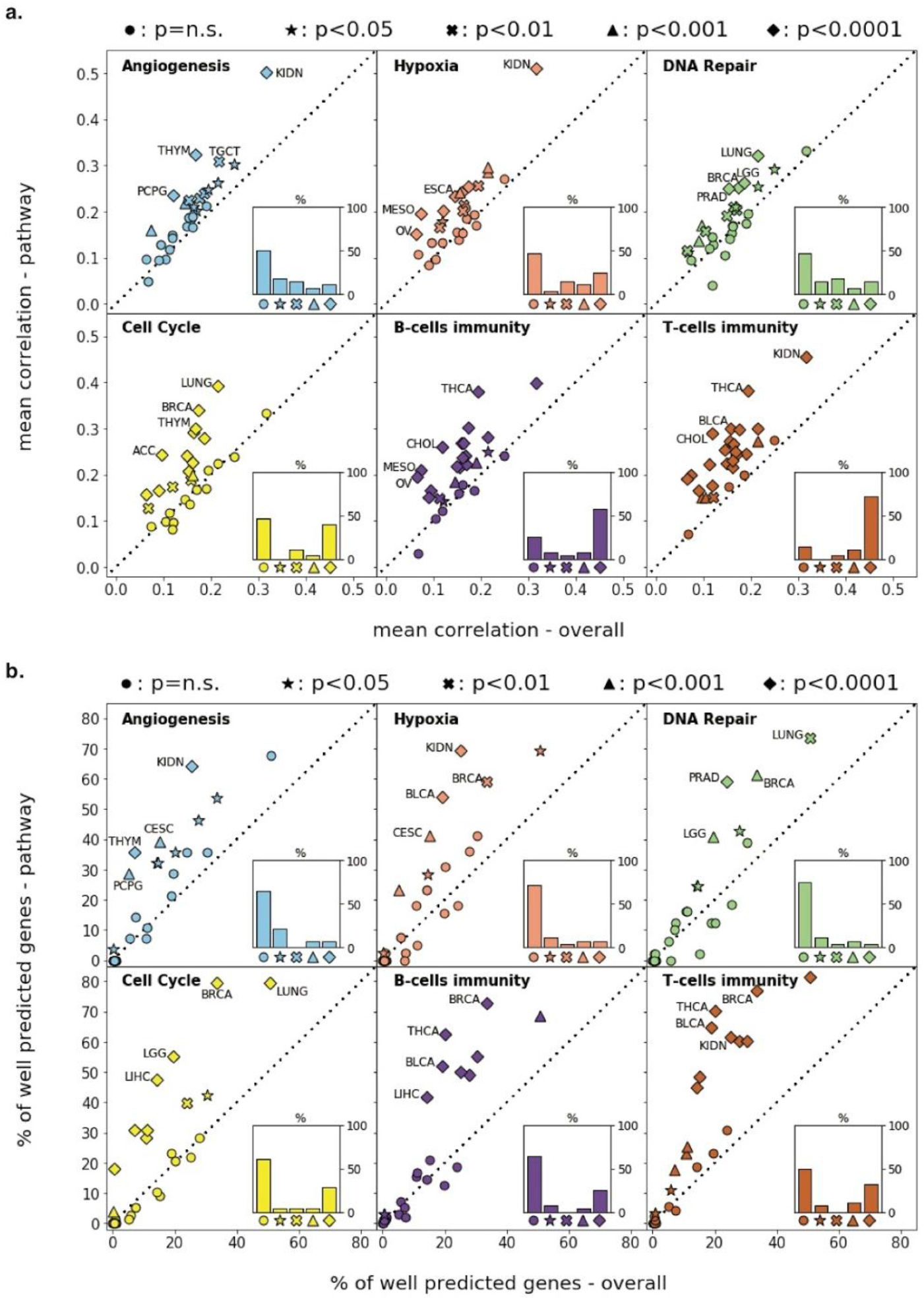
Prediction of signatures for cancer hallmarks. **a.** Comparison of correlation scores *R*_*p*_ for each gene pathway defined in Table S4 and involved in angiogenesis, hypoxia, DNA repair, cell cycle, and immune responses mediated by B and T cells, with the mean correlation coefficient *R*_*o*_ obtained for 10,000 random lists of the same number of genes, for all 28 cancer types from the TCGA dataset. The indicated statistical significance refers to the probability of obtaining a correlation *R* > *R*_*p*_ in the distribution of correlations for random lists, for each given cancer type. Insets show the percentages of the different cases of statistical significance between cancer types. The dotted line is the identity line *R*_*p*_ = *R*_*o*_. **b.** As in **a**, but in terms of percentage of genes considered well-predicted (as defined in the text and in Fig. 2).

As expected, the proportion of well-predicted genes from these six pathways increased with the size of the dataset for each cancer type (as in Fig. 2a). Indeed, the largest datasets (BRCA and LUNG) had higher proportions of genes accurately predicted per pathway and for the six defined pathways than smaller datasets (Supplementary Fig. S3). Nevertheless, HE2RNA predicted a significant proportion of gene expressions within these pathways in some of the smallest datasets: for example 18% of the cell cycle pathway genes were accurately predicted in pancreatic adenocarcinoma (PAAD), together with 29% of the angiogenesis network and 23% of the hypoxia pathway genes in pheochromocytoma and paraganglioma (PCPG). Moreover, these results confirmed our previous analysis (Fig. 2e) in which genes involved in cell cycle regulation were among the most accurately predicted for the BRCA dataset.

As a control experiment, we used HE2RNA to predict the levels of expression of housekeeping genes (HK, list in Supplementary Table 4). We did not expect the model to perform well on this subset of genes known to be expressed at similar levels in diverse cell types under normal and pathological conditions. As expected (Supplementary Fig. S4), in most cases, the predictions for this gene set were similar to those for randomly selected genes. Indeed, when the procedure described above was followed, the correlation for the HK signature was significantly better than that for a random set of genes in only 14% of cancer types, and this percentage dropped to zero for measurements of the proportion of well-predicted genes. This analysis validated our normalization procedure and the ability of HE2RNA to focus on specific cancer-related molecular information when trained on TCGA tissue images.

### HE2RNA, a tool for virtual spatialization

The HE2RNA model assigns a score to all super-tiles contributing to gene prediction and is therefore interpretable by design. Once a predictive model has been trained, it can identify the specific regions predictive of the expression level of a given gene on each WSI. The larger the number of super-tiles chosen for the model, the higher the definition of the spatialization will be. The limit is reached when all the tiles of the training WSIs are treated separately (*k*=8,000) (Supplementary Fig. S1). Previous studies^28,29,42^ have demonstrated that a virtual spatialization map (VSM), covering the entire WSI can be defined on the basis of CNN models. These heatmaps reflect the importance score assigned to each tile used in the algorithm. For validation of the accuracy of such VSMs, we considered the subset of genes encoding the human CD3 protein complex (*CD3D*, *CD3E*, *CD3G* and *CD247*)^45,46^, as an example of a well-predicted marker, for which we obtained a spatialization map (Fig. 4a). CD3 is a transmembrane receptor glycoprotein specifically expressed at the surface of T lymphocytes, involved in their activation^47^ and constituting an interesting biomarker of immune infiltration of the tumor microenvironment^48^. First, we applied the CD3 gene expression prediction model, trained as described above on TCGA, to an external H&E-stained image from a LIHC sample, to generate a full VSM. Our cross-validation procedure generated five models, each producing four predictions per tile (one for each gene). The expression levels of the four genes considered are strongly correlated (with inter-gene correlation coefficients exceeding 0.78 on TCGA-LIHC). We therefore averaged their expressions levels as well as the model predictions and obtained an overall mean correlation of *R*=0.43 for the TCGA-LIHC dataset. For the generation of the final VSM, we averaged the predictions obtained with the five models trained in cross-validation. We validated this VSM on the same LIHC sample that was washed and then stained with an anti-CD3 antibody. This procedure generated two WSIs with different staining patterns from the same slide (Fig. 4a-b). We calculated the correlation between the predicted mean activity per tile and the actual number of T cells obtained by using QuPath software on the CD3-stained slide. We obtained a correlation coefficient of *R*=0.54. Moreover, as HE2RNA focuses particularly on histological regions associated with higher levels of gene expression, we analyzed the 100 tiles for which the model predicted the highest value for the expression of CD3 genes. The median number of T cells in those tiles was 42.5 cells, whereas the median number of T cells on all 28,123 tiles of the slide was 4, confirming the accurate spatial interpretability of the predictive model (Fig. 4c). These results were also confirmed visually, by focusing on the tiles with the highest and lowest scores associated with CD3 gene expression (Fig. 4d-e).

**Fig. 4:**
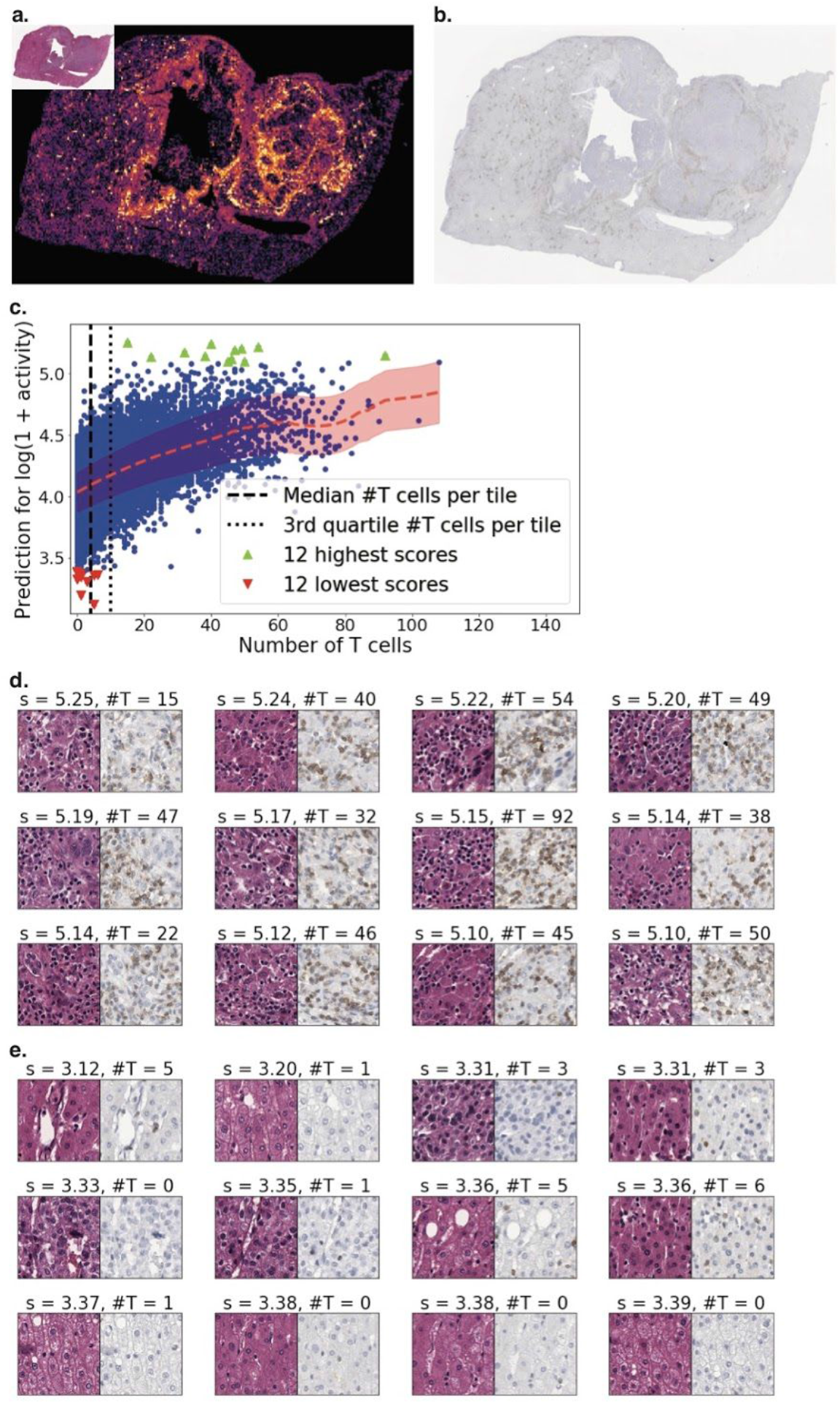
Virtual spatialization of *CD3* expression based on HE2RNA prediction, confirmed by CD3 immunohistochemistry. **a.** *Top left inset:* HE-stained slides were obtained for LIHC patients from a French hospital. *Main image:* Corresponding heatmap for the expression of CD3^+^-encoding genes expression predicted by our model. **b.** CD3 immunohistochemistry (IHC) results for the same slide were obtained by washing out H&E stain and staining the same slide for IHC. **c.** Pearson’s coefficient (*R*= 0.54) for the correlation between the percentage of cells with higher levels of CD3 expression predicted by our model and the percentage of CD3^+^ cells actually detected on the IHC slide. **d.** Extraction of the tiles associated with the highest score for CD3^+^-encoding genes expression predicted from the slide. S = Score corresponding to the log expression score of each tile; #T= Number of T cells per tile as determined with QuPath. **e.** As in **d** but for the tiles with the lowest score for CD3 expression.

Overall, these results demonstrate that our model can efficiently predict the spatial expression profile of a subset of genes. Such predictions are possible even for RNA-Seq data obtained for bulk cells rather than in single-cells transcriptomic approaches.

### HE2RNA for microsatellite instability status prediction

In addition to predicting gene expression and making it possible to generate VSMs, our HE2RNA model also provides a novel *transcriptomic representation* of the histological images, of potential utility for several different clinical and medical situations. In this approach, while learning the transcriptome, HE2RNA transformed each WSI into a vector of P features corresponding to the dimensionality of the last hidden representation of the neural network. We considered the other extreme of the preprocessing strategy (a single super-tile, corresponding to the mean of all tiles) and a last hidden layer of 256 neurons. As an example within the landscape of possibilities allowed by this new type of representation, we studied the problem of microsatellite Instability (MSI) status prediction directly from H&E-stained WSIs. DNA mismatch repair deficiency (MMRd) is most frequently observed in adrenocortical, rectal, colon, stomach, and endometrial tumors^49^, but it may also occur in cancers of the breast, prostate, bladder, and thyroid^50^. Tumors harboring this phenotype develop both point and frameshift mutations at an abnormally high rate and are often described as “hypermutated”^51^. The failure of mismatch repair to correct replication errors at tandem repeats of short DNA sequences known as microsatellites can lead to the phenomenon of high-level MSI (MSI-H)^51^. MSI-high status was recently identified as a predictor of the efficacy of anti-PD-1/PD-L1 immunotherapy^31,52^. MSI status can be determined by immunohistochemistry (staining for MLH1, MSH2, MSH6, PMS2) or genetic analyses (polymerase chain reaction (PCR)-based analysis of MSI markers)^53^. However, such screening is systematically performed only at high-volume tertiary care centers. There is, therefore, a great need to screen standard WSIs from patients with solid tumors with a high probability of MSI-H directly, to facilitate access to immunotherapy. A recent study^54^ showed that CNNs can learn to predict MSI status directly from histology slides for stomach adenocarcinoma and colorectal cancer. Based on these results, we collected, from the TCGA-COAD dataset, RNA-Seq measurements, WSIs and the corresponding MSI status of each patient from the dataset, to investigate the effects of integrating RNA-Seq information on the prediction of MSI status from pathology images. We divided the dataset in two classes, MSI-H and MSI-non high (MSI-NH), corresponding to both the MSI-Low and MSI-stable status subsets. In total, 441 patients matched these criteria, with a MSI-H/MSI-NH ratio of 0.18. We considered a setup in which the COAD dataset was randomly split between two different hospitals. In Hospital A, the first subset of WSIs was used to train a simplified version of HE2RNA (in which a mean over the tiles was also applied during the learning phase) to predict RNA-Seq data, but not MSI status (Fig. 5a). In Hospital B, the second subset of WSIs was used to train a binary classifier MSI-H vs. MSI-NH (without transcriptomic data). This use case corresponds to situations frequently encountered when a research center develops models for predicting treatment outcomes from a small dataset, with access to additional external data but not the corresponding biological status of interest. The approach proposed here could constitute a new paradigm in transfer learning in the field of medicine. First, to test the robustness of our model, we considered different relative size splits for the different data subsets used in the two hospitals, ranging from ⅙-⅚ (corresponding to 73 - 368 patients), to the inverse proportion. Moreover, each experiment, for a given ratio of Hospital A / Hospital B subset sizes, was repeated 50 times with different random splits, to generate robust performance estimates (Fig. 5b). For each split, we trained and compared two binary classifiers on the Hospital B dataset, each evaluated with a 10-times-bootstrapped 3-fold cross validation to predict MSI status. The first classifier consisted of a feed-forward neural network with two hidden layers (with 256 and 128 neurons, respectively) and was trained directly on WSIs from Hospital B. The second had a similar architecture, with a feed-forward neural network with one hidden layer (with 128 neurons), and was trained with the 256-dimensional transcriptomic representation learned from Hospital A and transferred to Hospital B WSIs. The performances of the two models are shown in Fig. 5b (AUC, area under the ROC curve) as a function of the proportion of samples in the two subsets. Two different regimes emerged. On one side (to the right of the figure), when only a few samples were used to learn the transcriptomic representation (Hospital A) and most patients were used to train the model for MSI prediction, the classifier trained directly on the WSIs (or more precisely on their 2048-dimensional ResNet representations) outperformed that based on the 256-dimensional transcriptomic representation of the WSI, learned through experience at Hospital A. At the other extreme, when only a few examples were available to train the classifiers at Hospital B (left side of the figure), the opposite pattern was observed. In this regime (in the example, for a split 3/4 - 1/4 of the samples between the two hospitals), the transcriptomic-based model gave more accurate predictions than the WSI-based model (two-tailed Wilcoxon test: *p*<0.0001; Fig. 5c). Finally, to confirm that the improvement in performances was not due purely to the dimensionality of the representation, but to the specific internal transcriptomic representation, we also considered the same MSI classifier trained with the 256-dimensional representation given by two different autoencoders, respectively trained on Hospital A and Hospital B subsets. A two-tailed Wilcoxon test confirmed (*p*<0.0001) that the classifier based on the transcriptomic representation of WSI slides outperformed that based on the other reduced dimension representations.

**Fig. 5:**
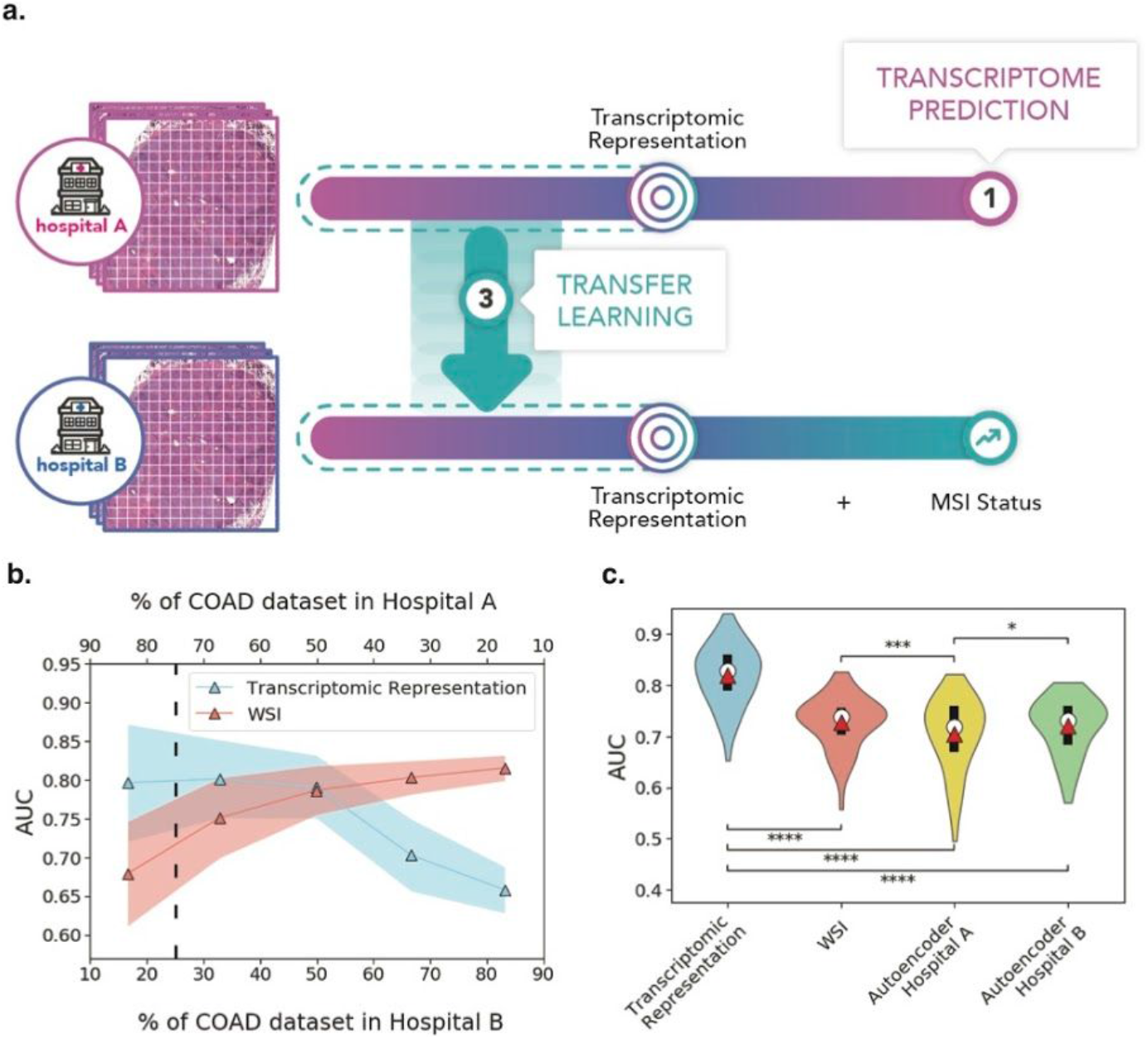
Prediction of microsatellite instability status using transfer learning from transcriptomic representation. **a.** Schematic diagram of the set-up studied. In Hospital A, a neural network is trained on COAD H&E slides for the prediction of gene expression. The internal lower-dimensional transcriptomic representation is then used in Hospital B to improve the prediction of microsatellite instability (MSI) status. **b**. Change in area under the ROC curve (AUC) (solid lines: mean, shaded area: 68% confidence interval) for the model based on the transcriptomic representation learned in Hospital A and trained on Hospital B (blue) and the model directly based on WSI images from Hospital B (red), as a function of the fraction of the COAD subset used in the two hospitals. **c**. Violin plots for the distribution of AUC values for the neural network MSI status classifier at Hospital B, trained on the 256-dimensional transcriptomic representation, WSI images, and a 256-dimensional representation given by two different autoencoders, respectively, trained on the hospital A and B subsets. Significance for the two-tailed Wilcoxon test: * (*p* <0.05), *** (*p* <0.001) and **** (*p* <0.0001). For each case, we indicate the median (white dot), mean (red triangle), and quartiles (black bar). WSI: whole-slide image. COAD: colorectal adenocarcinoma.

## Discussion

Our results demonstrate that CNNs, such as HE2RNA, can be used to predict the expression of a large subset of coding and non-coding genes, in various cancer types, after training on the corresponding histological images. They confirm that the transcriptomic information is actually encoded in tissue biopsy specimens and that a deep learning model can learn and predict it. HE2RNA robustly and consistently predicted subsets of genes as expressed in different cancer types, including genes involved in immune cell activation status and signaling in particular. One possible reason for this is that the algorithm can recognize immune cells and correlate their presence with the expression of a subset of protein-coding genes (such as *CQ1B*). Major breakthroughs in cancer therapy are driven by discoveries of treatment targets for immunotherapy in many types of cancer. Such machine-learning tools could, therefore, be used to predict immune infiltration in some tumor types on the basis of biopsy slides only, without the need for next-generation sequencing, which would help to broaden access to these new therapies. In the future, it would be interesting to determine whether similar models could be trained to predict patient response to immunotherapy, making it possible to identify histology-based biomarkers of treatment response.

We have shown that HE2RNA can also correctly predict the expression of genes involved in more cancer type-specific pathways, such as fibrosis in hepatocellular carcinoma, or *CHK* gene expression in breast cancers. The accuracy of our model for detecting biological changes, and molecular and cellular modifications within cancer cells, was also confirmed by the greater prediction accuracy for defined gene signatures than for lists of random genes. In the last decade, studies have increasingly shown that biopsy specimens and tissue sections contain tremendous amounts of information^55,56,57^. Our model might be able to recognize more subtle structures in the tissue images and more interesting histological patterns might emerge, shedding light on the specific tumoral regions important for the development of specific cancer types. Strikingly, even with RNA-Seq profiles obtained from bulk cells isolated from whole histological slides rather than sorted single cells, our model was still able to predict the spatial concentration of CD3 expression from the H&E-stained slide alone. Such methods could be extended to other genes, including genes related to immune activity in particular, the expression levels of which were well predicted by our model, and which could represent a major tool for medical diagnosis and prediction, by providing virtual multiplexed staining for all samples stained with H&E alone. These new approaches may help to overcome technical issues in IHC, such as fixation or antigen retrieval, together with the high level of variability between observers.

CNNs for image recognition make use of an internal representation of the original data that they infer. The features of this latent space encode the statistics of natural images and the information of importance for image recognition. Similarly, the internal transcriptomic representation, learned by HE2RNA during the prediction of RNA-Seq data, may constitute an important step towards understanding the biological descriptors required for medical/clinical classification problems and the link between the information contained at the tissue and molecular levels. Finally, we have shown that the lower-dimensional transcriptomic representation learned during the RNA-Seq prediction task can be very powerful when transferred to other datasets used for a different task, as frequently observed when deep learning is applied to digital pathology (i.e. the use of transfer learning from CNN trained on ImageNet datasets to questions related to histological images, such as prediction of patient survival or response to treatment). This seems to be particularly true for small WSI datasets, for which even partial information about the connection between histological and molecular information can significantly improve the performance of deep learning models. We used MSI status prediction from a small dataset of H&E-stained images as a representative case study. We showed that the use of a transcriptomic representation in a transfer learning framework outperformed similar models based on less informative representations, such as WSI only. It has recently been shown that the MSI status of patients affects their response to immunotherapy^31,52^. Our HE2RNA model could therefore be used in the future to facilitate the definition of patient MSI status and to provide easier access to immunotherapies for a larger number of eligible patients. The practical implications of predicting gene expression level from H&E slides should not be underestimated. In the future, the performance of this model may improve considerably, through the use of larger, richer datasets for training, and this model could also be used in different scenarios, to determine gene signatures or immune infiltration on the basis of histologic slides only. It would therefore constitute an interesting tool for enabling patients to gain access to personalized medicine even in hospitals that do not yet have access to next-generation sequencing technologies.

## Acknowledgements

We thank The Cancer Genome Atlas (TCGA) program for providing free access to histology slide images and RNA-Seq data, and Qiagen Bioinformatics for allowing us to use their Ingenuity Pathway Analysis (IPA) software on free trial. We thank Simon Jégou, Aurélie Kamoun, Charles Maussionet, Eric Tramel and for insightful discussions.

## Author contributions

B.S., J.C., M.S., M.Z., T.C., P.C. and G.W. designed the experiments; P.M. performed the double-staining experiments; B.S., A.R., C.S. and P.C. performed the numerical experiments and wrote the code to achieve the different tasks; B.S., A.R., E.P., M.M. and P.C., analyzed the results. B.S., A.R., E.P., S.T., M.M., P.C. and G.W. wrote the manuscript with the assistance and feedback of all the other co-authors.

## Methods

### TCGA Pan-Cancer Dataset

This study was based on publicly available data from TCGA. We selected samples from primary tumors only, for which both RNA-Seq and WSI data were available. Transcriptomic data (FPKM-UQ) were extracted from frozen tissues, and the slides analyzed were digitized HE-stained formalin-fixed, paraffin-embedded (FFPE) histology slides, referred to here as whole-slide images (WSIs). WSIs were available for the following cancer types (and corresponding abbreviations):

**Table 1:** TCGA dataset, detailed information. A matched WSI RNA-Seq data pair is considered here to be a sample.

| Project ID |  | #Patients | # Samples |
| --- | --- | --- | --- |
| BRCA | Invasive breast carcinoma | 1057 | 1131 |
| LUNG | Lung adenocarcinoma (LUAD) | 944 | 1046 |
|  | Lung squamous cell carcinoma (LUSC) |  |  |
| KIDN | Kidney chromophobe carcinoma (KICH) | 843 | 882 |
|  | Kidney renal clear cell carcinoma (KIRC) |  |  |
|  | Kidney renal papillary cell carcinoma (KIRP) |  |  |
| LGG | Brain lower-grade glioma | 485 | 843 |
| SARC | Sarcoma | 252 | 597 |
| UTER | Uterine carcinosarcoma (UCS) | 558 | 673 |
|  | Uterine corpus endometrial carcinoma (UCEC) |  |  |
| THCA | Thyroid carcinoma | 497 | 509 |
| COAD | Colon adenocarcinoma | 445 | 463 |
| HNSC | Head and neck squamous cell carcinoma | 423 | 462 |
| BLCA | Bladder urothelial carcinoma | 383 | 456 |
| PRAD | Prostate adenocarcinoma | 399 | 451 |
| LIHC | Liver hepatocellular carcinoma | 359 | 373 |
| STAD | Stomach adenocarcinoma | 350 | 371 |
| CESC | Cervical squamous cell carcinoma and endocervical adenocarcinoma | 267 | 279 |
| TGCT | Testicular germ cell tumors | 149 | 245 |
| ACC | Adrenocortical carcinoma | 55 | 224 |
| GBM | Glioblastoma multiforme | 96 | 212 |
| PAAD | Pancreatic adenocarcinoma | 175 | 195 |
| PCPG | Pheochromocytoma and paraganglioma | 175 | 194 |
| THYM | Thymoma | 116 | 176 |
| READ | Rectum adenocarcinoma | 159 | 160 |
| ESCA | Esophageal carcinoma | 133 | 135 |
| SKCM | Skin, cutaneous melanoma | 97 | 108 |
| MESO | Mesothelioma | 74 | 94 |
| UVM | Uveal melanoma | 80 | 80 |
| OV | Ovarian serous cystadenocarcinoma | 74 | 75 |
| DLBC | Lymphoid neoplasm diffuse large B cell lymphoma | 44 | 44 |
| CHOL | Cholangiocarcinoma | 36 | 36 |

### Cleaning

Gene expression data were available for 60,483 fragments, many corresponding to non-coding genomic regions. We chose to exclude genes with a median expression of zero (i.e. not expressed in more than half the samples considered), to improve the interpretability of the results. After the application of this filter, 30,839 genes remained, 17,759 of which encoded proteins (all Ensembl genes associated with a corresponding Ensembl protein ID, for the Hg19 human genome sequence). Our selection included almost 90% of known protein-coding genes.

### Preprocessing

The application of deep learning algorithms to histological data is a challenging problem, particularly due to the high dimensionality of the data (up to 100,000 × 100,000 pixels for a single whole-slide image) and the small size of available datasets. We divided the whole-slide images into squares of 112 × 112 μm (224 × 224 pixels) called “tiles”, and used the Otsu algorithm to select only those containing tissue, excluding the white background. We sampled a maximum of 8,000 such tiles from each slide. We then extracted 2,048 features from those tiles with a 50-layer ResNet pretrained on the ImageNet dataset, such that a slide could be represented as a 8,000 × 2,048 matrix.

For the first phase of this work (transcriptome prediction), we accelerated the training of our models through a simple preprocessing step inspired by simple linear iterative clustering (SLIC)^58^: we used the k-means algorithm to create 100 clusters (super-tiles) of tiles on the basis of tile location on the slide, and we averaged the features of the tiles within each cluster. The use of these super-tiles, reduces the dimensions of a slide to 100 × 2,048. The model was first trained on this reduced dataset, with all the TCGA data. Then, for specific organs, fine-tuning was achieved with full-scale data from the organs concerned only.

Gene expressions values covered several orders of magnitude. Thus, regression analysis directly on raw RNA-Seq data would lead the model to focus only on the most strongly expressed genes, which would dominate the mean squared error. We overcame this problem by an a → log_10_(1 + a) transformation on gene expressions.

### Model architecture

The HE2RNA model is a multilayer perceptron (MLP), applied to every tile (or super-tile) of the slide. This choice, as opposed to a simple linear regression, allows to perform multitask learning by taking into account the correlations between multiple gene expressions at the level of (super-)tiles. For practical purposes, this is equivalent to applying successive 1D convolutions of kernel size 1 and stride 1 to slide data. The activation function is a rectified linear unit and dropout is applied between consecutive layers. For an input matrix of size n_tiles_ × 2,048 (with n_tiles_ = 100 or n_tiles_ = 8,000), the output of this neural network is a matrix of size n_tiles_ × n_genes_, where n_genes_ is the number of genes for which expression is to be predicted. Thus, the model produces one prediction per gene and per tile, but the real value is available only at the scale of the slide. For this reason, tile predictions must be aggregated for model training and the calculation of metrics.

During the training phase, the model selects a number *k* at every iteration and for every gene, and averages only the *k* highest tile predictions, to produce the slide-level prediction. Indeed, we are predicting the logarithm of gene expression, and the (super-)tiles with the highest level of expression should contribute the most to this value. The number *k* is sampled from the list (1, 2, 5, 10, 20, 50, 100) for super-tile-preprocessed data, and from the list (10, 20, 50, 100, 200, 500, 1000, 2000, 5000) for full-scale data. This increases the difficulty of the task and thus reduces overfitting. During inference, slide-level predictions obtained with every possible value of *k* are averaged. This is equivalent to calculating a weighted mean of per-tile predictions for every gene, with a greater weight for tiles for which the model predicts high levels of expression.

### Training and evaluation

HE2RNA was trained with a five-fold cross-validation designed to meet the following requirements: every sample from a patient should be in the same fold, and, when training on all TCGA data, TCGA projects should be evenly distributed between folds. When training a model on a single subset (e.g. BRCA), we ensured that the folds used for cross-validation were consistent with those used for all the TCGA data. The model simultaneously predicting all genes for all types of cancers was trained on all TCGA data (10514 samples), with super-tile preprocessing.

When specific sets of genes and organs (such as CD3 in the liver) were used to produce a precise heatmap of gene expression, the model was trained at the finest scale.

To optimize the trade-off between achieving optimal performance using full-scale TCGA data (10,514 slides × 8,000 tiles × 2,048 features) and minimizing the machine time for training, we first trained the model for 200 epochs on all SLIC-preprocessed TGCA data (10,514 slides × 100 tiles × 2,048 features), before fine-tuning it for 50 epochs on full-scale data for the organ of interest (373 slides × 8,000 tiles × 2,048 features for LIHC).

The performance metric was always calculated separately for each organ considered, to prevent bias. Gene expression levels can differ considerably between organs and a good performance for gene expression could be achieved simply through recognition of the organ of origin.

### GSEA signatures

Using Gene Set Enrichment Analysis (GSEA) software and Molecular Signatures Database v6.2, we obtained the following signatures for the following pathways. For each pathway we then selected the genes present in at least two of the chosen signatures. (See Table S4 for a complete list of the genes retained for each pathway).

**Table 2:** List of signatures from GSEA combined to define the six cancer pathways.

|  |  |
| --- | --- |
| **Angiogenesis** | HALLMARK_ANGIOGENESIS,<br>BIOCARTA_VEGF_PATHWAY,<br>KEGG_VEGF_SIGNALING_PATHWAY,<br>GO_ANGIOGENESIS |
| **Hypoxia** | HALLMARK_HYPOXIA,<br>BIOCARTA_HIF_PATHWAY,<br>GO_REGULATION_OF_CELLULAR_RESP<br>ONSE_<br>TO_HYPOXIA |
| **DNA repair** | HALLMARK_DNA_REPAIR,<br>REACTOME_DNA_REPAIR,<br>GO_DNA_REPAIR |
| **Cell cycle** | BIOCARTA_CELLCYCLE_PATHWAY,<br>KEGG_CELL_CYCLE,<br>REACTOME_CELL_CYCLE,<br>GO_CELL_CYCLE |
| **B cell-mediated immunity** | GO_B_CELL_MEDIATED_IMMUNITY |
| **T cell-mediated immunity** | REACTOME_ADAPTATIVE_IMMUNE_<br>SYSTEM<br>GO_ADAPTATIVE_IMMUNE_RESPONSE<br>GO_REGULATION_OF_ADAPTATIVE_IMM<br>UNE_RESPONSE |

### Visual spatialization and double staining

Features were extracted from every tile containing matter, with the ResNet50 algorithm. The model was then used to calculate a score for each tile. Finally, the heatmap at the scale of the whole slide was obtained by weighting each tile by its score. For instance, for the double-stained HE-CD3 slide, we extracted a total of 23,123 tiles to generate the heatmap shown in Fig. 3.

The tissue section was first stained with hematein, eosin and saffron, coverslipped and scanned using a Leica Aperio Scanner. The coverslip and the mounting reagents were removed using acetone, and the slide was further unstained using an alcohol/acid solution (1%). Immunostaining was performed using a Leica Bondmax Autostainer (Leica Biosystems, Wetzlar, Germany) with a anti-CD3 antibody (Dako, Santa Clara, California, Clone, F7.2.38, dilution 1/50), according to manufacturer's instructions. Antigen retrieval was performed with E2 reagent, and Immunodetection was performed using Polymer and 3,3’ di-aminobenzidine (Bond Polymer Refine Detection Kit, Leica Biosystems).The immunostained slide was further scanned.

For comparison of the predicted heat-map and the CD3-stained WSI, we used QuPath software on the WSIs to estimate the actual number of T cells per tile, and then calculated the correlation between this number and the score per tile.

### MSI status and patient cohort

We used the histology images for *n* = 441 patients with colorectal carcinoma (COAD) (diagnostic slides, FFPE tissue) from the TCGA dataset, together with the corresponding MSI status data. For transcriptomic learning at Hospital A, we used the simplest version of HE2RNA, in which the input for each slide was the mean value over the ResNet50 representation of every tile (see Fig. S1).

We also set the MLP architecture as follows: two hidden dense layers of 1024 and 256 neurons with sigmoid activation followed by the last prediction layer with 28334 output neurons and linear activation (28334 being the number of coding or non-coding genes with non-zero median expression levels over the samples of COAD dataset). The model was trained for 50 epochs without hyperparameter tuning. For the MSI classifier at Hospital B, we compared two different models. The first one was an MLP with two hidden dense layers of 256 and 128 neurons and a one-neuron output layer, all with sigmoid activation, also fed with the all-image average of the ResNet50 representation of the tiles and trained for 50 epochs. The second consisted of a neural network with the same architecture, without the first hidden layer of 256 neurons; this second classifier was trained on the 256-dimensional representations of Hospital B WSIs, given by the transcriptomic representation learned at Hospital A, or by an autoencoder (with a mirror architecture and three hidden dense layers of 1024, 256 and 1024 neurons, with ReLU and linear activations), trained on either the Hospital A or Hospital B subset. We performed different experiments, for different ratios of sample size between the Hospital A and B subsets. We performed a three-fold cross-validation on the Hospital A task, and the transcriptomic representation for the Hospital B dataset was obtained by averaging the three corresponding inferences for the Hospital B subset. MSI status was also predicted in a three-fold cross-validation setup. Moreover, for each A-/B-subset size ratio, the random split between the two hospitals was bootstrapped 50 times to generate robust performance estimates.

### Statistics and reproducibility

We determined whether the correlation between the prediction of the RNA-Seq expression levels for a given gene and the real values was statistically significant, by comparison with the distribution of correlations predicted by a model with the same architecture as HE2RNA but untrained. The estimated *p*-values for tests against the null hypothesis of random correlations were then corrected by the Holm-Šidák method, to account for multiple comparisons. A gene was considered significantly well predicted, for a given cancer type, if its corrected p-value was below 0.05. The IPA analysis in Fig. 1c, 1d and 1e is based on Fisher’s exact tests. The direction of the change in gene expression is not taken into account in this calculation. For the analysis on LIHC and BRCA, given the large number of genes with *p*-value below 0.05, we focused on genes for which the coefficient of the correlation between the predicted and true value was greater than 0.4: 786 genes for BRCA and 765 for LIHC.

For the analysis of the hallmark of cancer pathways, we selected, for each pathway and cancer type, 10,000 different random lists of genes of the same length as the pathway list. We then determined the correlation for all genes and the number of genes for which expression was well predicted (in terms of the Holm-Šidák corrected *p*-value), for all these lists.

We compared the mean correlation *R*_*p*_ over the pathway gene list, with the distribution of the mean correlation over the random gene lists. We plotted *R*_*p*_ against the mean *R*_*o*_ over all the 10,000 averages of *R*_*r*_ for the random list (Fig. 3). We considered a given pathway for a given cancer type to be better predicted than a corresponding random set, when the probability *p* of *R*_*r*_ > *R*_*p*_ was <0.05.

We adopted the same approach for the percentage of well predicted genes as a measure, but compared the percentages *f*_*p*_ of well-predicted genes in the pathway, with the distribution *f*_*o*_ over the 10,000 random lists.

The MSI prediction scores are expressed as area under the ROC curve (AUC) and a two-tailed Wilcoxon test was used to compare the different distributions of scores.

### Data availability

The TCGA dataset is publicly available via the TCGA portal (https://portal.gdc.cancer.gov). All other relevant data are present in the article or supplementary files, or are available from the author upon request.

### Code availability

The code used for training the models has a large number of dependencies on internal tooling and its release is therefore not feasible. However, all experiments and implementation details are described thoroughly in the Methods so that it can be independently replicated with non-proprietary libraries.

## References

1. Zarella, M. D. et al. A Practical Guide to Whole Slide Imaging: A White Paper From the Digital Pathology Association. Arch. Pathol. Lab. Med. 143, 222–234 (2019).

2. Mukhopadhyay, S. et al. Whole Slide Imaging Versus Microscopy for Primary Diagnosis in Surgical Pathology: A Multicenter Blinded Randomized Noninferiority Study of 1992 Cases (Pivotal Study). Am. J. Surg. Pathol. 42, 39–52 (2018).

3. Wang, H. et al. Mitosis detection in breast cancer pathology images by combining handcrafted and convolutional neural network features. J. Med. Imaging Bellingham Wash 1, 034003 (2014).

4. Cireşan, D. C., Giusti, A., Gambardella, L. M. & Schmidhuber, J. Mitosis detection in breast cancer histology images with deep neural networks. Med. Image Comput. Comput.-Assist. Interv. MICCAI Int. Conf. Med. Image Comput. Comput.-Assist. Interv. 16, 411–418 (2013).

5. Turkki, R., Linder, N., Kovanen, P. E., Pellinen, T. & Lundin, J. Antibody-supervised deep learning for quantification of tumor-infiltrating immune cells in hematoxylin and eosin stained breast cancer samples. J. Pathol. Inform. 7, 38 (2016).

6. Esteva, A. et al. Dermatologist-level classification of skin cancer with deep neural networks. Nature 542, 115–118 (2017).

7. Hou, L. et al. Patch-based Convolutional Neural Network for Whole Slide Tissue Image Classification. Proc. IEEE Comput. Soc. Conf. Comput. Vis. Pattern Recognit. 2016, 2424–2433 (2016).

8. Khoshdeli, M., Borowsky, A. & Parvin, B. Deep Learning Models Differentiate Tumor Grades from H&E Stained Histology Sections. Conf. Proc. Annu. Int. Conf. IEEE Eng. Med. Biol. Soc. IEEE Eng. Med. Biol. Soc. Annu. Conf. 2018, 620–623 (2018).

9. Bulten, W. et al. Automated Gleason Grading of Prostate Biopsies using Deep Learning. ArXiv190707980 Cs Eess (2019).

10. Coudray, N. et al. Classification and mutation prediction from non-small cell lung cancer histopathology images using deep learning. Nat. Med. 24, 1559–1567 (2018).

11. Mobadersany, P. et al. Predicting cancer outcomes from histology and genomics using convolutional networks. Proc. Natl. Acad. Sci. U. S. A. 115, E2970–E2979 (2018).

12. Montalto, M. C. & Edwards, R. And They Said It Couldn’t Be Done: Predicting Known Driver Mutations From H&E Slides. J. Pathol. Inform. 10, 17 (2019).

13. Vamathevan, J. et al. Applications of machine learning in drug discovery and development. Nat. Rev. Drug Discov. 18, 463–477 (2019).

14. Schaumberg, A. J., Rubin, M. A. & Fuchs, T. J. H&E-stained Whole Slide Image Deep Learning Predicts SPOP Mutation State in Prostate Cancer. (Pathology, 2016). doi:10.1101/064279

15. Chang, P. et al. Deep-Learning Convolutional Neural Networks Accurately Classify Genetic Mutations in Gliomas. Am. J. Neuroradiol. 39, 1201–1207 (2018).

16. Kim, R. H. et al. A Deep Learning Approach for Rapid Mutational Screening in Melanoma. (Pathology, 2019). doi:10.1101/610311

17. Noorbakhsh, J. et al. Pan-cancer classifications of tumor histological images using deep learning. (Bioinformatics, 2019). doi:10.1101/715656

18. Xu, H., Park, S., Lee, S. H. & Hwang, T. H. Using transfer learning on whole slide images to predict tumor mutational burden in bladder cancer patients. (Bioinformatics, 2019). doi:10.1101/554527

19. Segal, E., Friedman, N., Kaminski, N., Regev, A. & Koller, D. From signatures to models: understanding cancer using microarrays. Nat. Genet. 37 Suppl, S38–45 (2005).

20. Lander, E. S. Array of hope. Nat. Genet. 21, 3–4 (1999).

21. Serratì, S. et al. Next-generation sequencing: advances and applications in cancer diagnosis. OncoTargets Ther. 9, 7355–7365 (2016).

22. Stratton, M. R., Campbell, P. J. & Futreal, P. A. The cancer genome. Nature 458, 719–724 (2009).

23. Conesa, A. et al. A survey of best practices for RNA-seq data analysis. Genome Biol. 17, 13 (2016).

24. Anders, S. & Huber, W. Differential expression analysis for sequence count data. Genome Biol. 11, R106 (2010).

25. Mortazavi, A., Williams, B. A., McCue, K., Schaeffer, L. & Wold, B. Mapping and quantifying mammalian transcriptomes by RNA-Seq. Nat. Methods 5, 621–628 (2008).

26. Kamps, R. et al. Next-Generation Sequencing in Oncology: Genetic Diagnosis, Risk Prediction and Cancer Classification. Int. J. Mol. Sci. 18, (2017).

27. McDermott, U., Downing, J. R. & Stratton, M. R. Genomics and the continuum of cancer care. N. Engl. J. Med. 364, 340–350 (2011).

28. Rivenson, Y. et al. Virtual histological staining of unlabelled tissue-autofluorescence images via deep learning. Nat. Biomed. Eng. 3, 466–477 (2019).

29. Saltz, J. et al. Spatial Organization and Molecular Correlation of Tumor-Infiltrating Lymphocytes Using Deep Learning on Pathology Images. Cell Rep. 23, 181–193.e7 (2018).

30. Shamai, G. et al. Artificial Intelligence Algorithms to Assess Hormonal Status From Tissue Microarrays in Patients With Breast Cancer. JAMA Netw. Open 2, e197700 (2019).

31. Le, D. T. et al. Mismatch repair deficiency predicts response of solid tumors to PD-1 blockade. Science 357, 409–413 (2017).

32. Merienne, N. et al. Cell-Type-Specific Gene Expression Profiling in Adult Mouse Brain Reveals Normal and Disease-State Signatures. Cell Rep. 26, 2477–2493.e9 (2019).

33. Nassiri, I. & McCall, M. N. Systematic exploration of cell morphological phenotypes associated with a transcriptomic query. Nucleic Acids Res. 46, e116–e116 (2018).

34. Courtiol, P., Tramel, E. W., Sanselme, M. & Wainrib, G. Classification and Disease Localization in Histopathology Using Only Global Labels: A Weakly-Supervised Approach. ArXiv180202212 Cs Stat (2018).

35. Love, M. I., Huber, W. & Anders, S. Moderated estimation of fold change and dispersion for RNA-seq data with DESeq2. Genome Biol. 15, 550 (2014).

36. Kleczko, E. K., Kwak, J. W., Schenk, E. L. & Nemenoff, R. A. Targeting the Complement Pathway as a Therapeutic Strategy in Lung Cancer. Front. Immunol. 10, 954 (2019).

37. Todros-Dawda, I., Kveberg, L., Vaage, J. T. & Inngjerdingen, M. The tetraspanin CD53 modulates responses from activating NK cell receptors, promoting LFA-1 activation and dampening NK cell effector functions. PloS One 9, e97844 (2014).

38. Medley, Q. G. et al. Characterization of GMP-17, a granule membrane protein that moves to the plasma membrane of natural killer cells following target cell recognition. Proc. Natl. Acad. Sci. U. S. A. 93, 685–689 (1996).

39. Sakurai, T. & Kudo, M. Molecular Link between Liver Fibrosis and Hepatocellular Carcinoma. Liver Cancer 2, 365–366 (2013).

40. Apostolou, P. & Papasotiriou, I. Current perspectives on CHEK2 mutations in breast cancer. Breast Cancer Dove Med. Press 9, 331–335 (2017).

41. Zrihan-Licht, S. et al. Association of csk-homologous kinase (CHK) (formerly MATK) with HER-2/ErbB-2 in breast cancer cells. J. Biol. Chem. 272, 1856–1863 (1997).

42. Sutherland, R. L. & Musgrove, E. A. Cyclins and breast cancer. J. Mammary Gland Biol. Neoplasia 9, 95–104 (2004).

43. Hanahan, D. & Weinberg, R. A. The hallmarks of cancer. Cell 100, 57–70 (2000).

44. Hanahan, D. & Weinberg, R. A. Hallmarks of cancer: the next generation. Cell 144, 646–674 (2011).

45. Tunnacliffe, A., Buluwela, L. & Rabbitts, T. H. Physical linkage of three CD3 genes on human chromosome 11. EMBO J. 6, 2953–2957 (1987).

46. Loh, E. Y. et al. Identification and sequence of a fourth human T cell antigen receptor chain. Nature 330, 569–572 (1987).

47. Smith-Garvin, J. E., Koretzky, G. A. & Jordan, M. S. T cell activation. Annu. Rev. Immunol. 27, 591–619 (2009).

48. Schreiber, R. D., Old, L. J. & Smyth, M. J. Cancer immunoediting: integrating immunity’s roles in cancer suppression and promotion. Science 331, 1565–1570 (2011).

49. Cortes-Ciriano, I., Lee, S., Park, W.-Y., Kim, T.-M. & Park, P. J. A molecular portrait of microsatellite instability across multiple cancers. Nat. Commun. 8, 15180 (2017).

50. Win, A. K. et al. Colorectal and other cancer risks for carriers and noncarriers from families with a DNA mismatch repair gene mutation: a prospective cohort study. J. Clin. Oncol. Off. J. Am. Soc. Clin. Oncol. 30, 958–964 (2012).

51. Boland, C. R. & Goel, A. Microsatellite instability in colorectal cancer. Gastroenterology 138, 2073–2087.e3 (2010).

52. Lemery, S., Keegan, P. & Pazdur, R. First FDA Approval Agnostic of Cancer Site - When a Biomarker Defines the Indication. N. Engl. J. Med. 377, 1409–1412 (2017).

53. Kather, J. N., Halama, N. & Jaeger, D. Genomics and emerging biomarkers for immunotherapy of colorectal cancer. Semin. Cancer Biol. 52, 189–197 (2018).

54. Kather, J. N. et al. Deep learning can predict microsatellite instability directly from histology in gastrointestinal cancer. Nat. Med. 25, 1054–1056 (2019).

55. Beck, A. H. et al. Systematic analysis of breast cancer morphology uncovers stromal features associated with survival. Sci. Transl. Med. 3, 108ra113 (2011).

56. Whitney, J. et al. Quantitative nuclear histomorphometry predicts oncotype DX risk categories for early stage ER+ breast cancer. BMC Cancer 18, 610 (2018).

57. Rawat, R. R., Ruderman, D., Macklin, P., Rimm, D. L. & Agus, D. B. Correlating nuclear morphometric patterns with estrogen receptor status in breast cancer pathologic specimens. NPJ Breast Cancer 4, 32 (2018).

## References

58. R. Achanta, A. Shaji, K. Smith, A. Lucchi, P. Fua, S. Susstrünk. SLIC Superpixels Compared to State-of-the-art Superpixel Methods, IEEE Transactions on Pattern Analysis and Machine Intelligence, vol. 34, No. 11, 2274 - 2282 (2022).

