## Supplemental figures and tables for "Transcriptomic learning for digital pathology"

### Supplementary Information

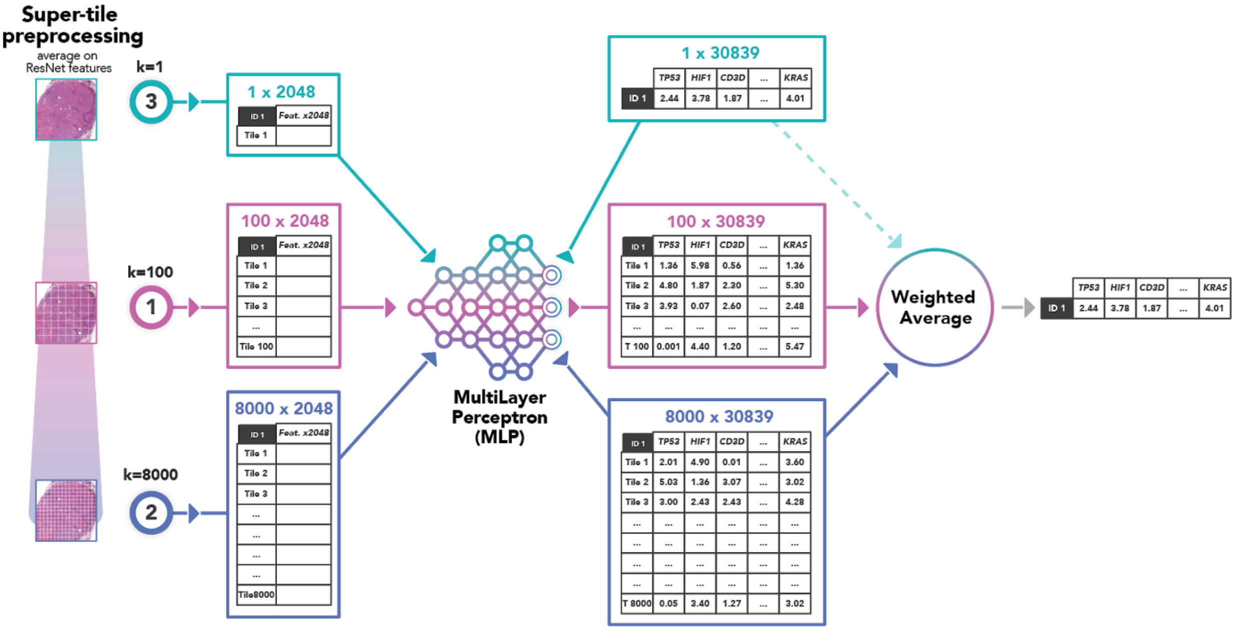

**Figure S1. Preprocessing and model structure.** A preprocessing algorithm, inspired by simple linear iterative clustering (SLIC) is applied to the 8,000 tiles into which the whole-slide images of the training set are divided, to produce *k* super-tiles. An average is obtained for each super-tile, at the level of the 2,048 features extracted from each tile using a 50-layer ResNet pretrained on the ImageNet. The number of clusters is decided according to the task (color and numerical code as in Fig. 1). A multilayer perceptron is applied to each cluster of the slide. The last layer of the model encodes the transcriptomic representation described in the text. This representation is then used to produce a prediction per cluster and per Ensembl gene of the RNA-Seq dataset. Finally, a weighted average (described in the Methods) provides the output prediction of gene expression associated with the slide.

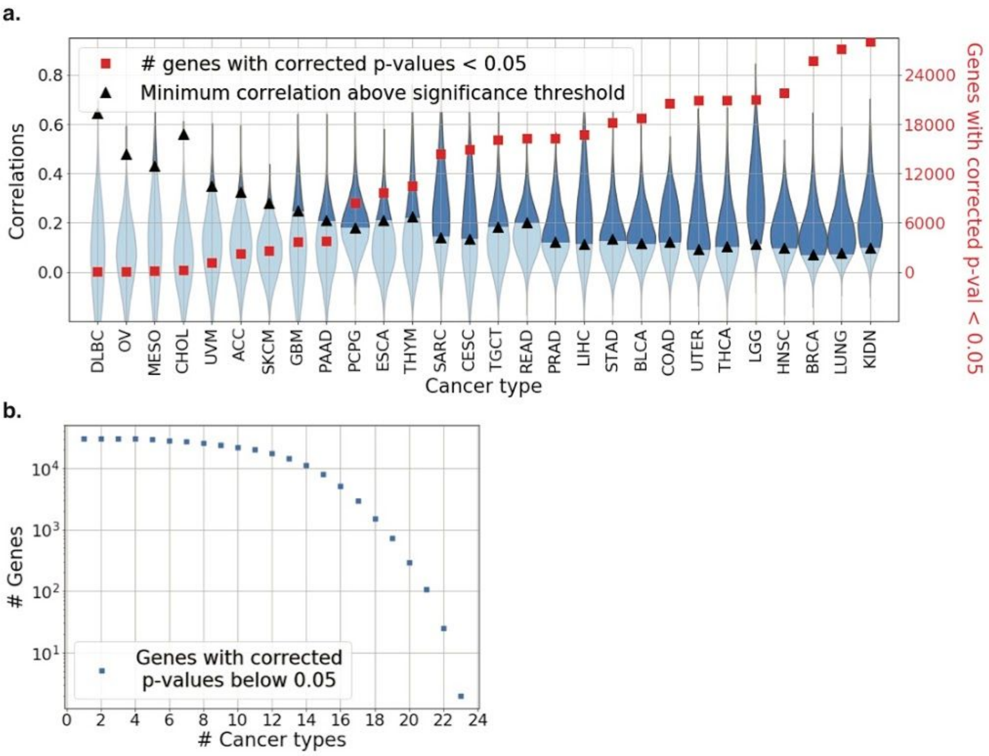

**Figure S2. Predictions of gene expression after Benjamini–Hochberg correction for the testing of multiple hypotheses. a**. Distribution of Pearson's correlation coefficients *R* (left axis, blue violin plots) and the number of coding and non-coding genes (right axis, red squares) with Benjamini–Hochberg-corrected *p*-values <0.05, for twenty eight cancer types from the TCGA. Black triangles indicate the minimal correlation coefficient required for significance in any given dataset. **b.** Number of coding and non-coding genes for which expression was significantly well-predicted for a given number of cancer types, as a function of the number of cancers.

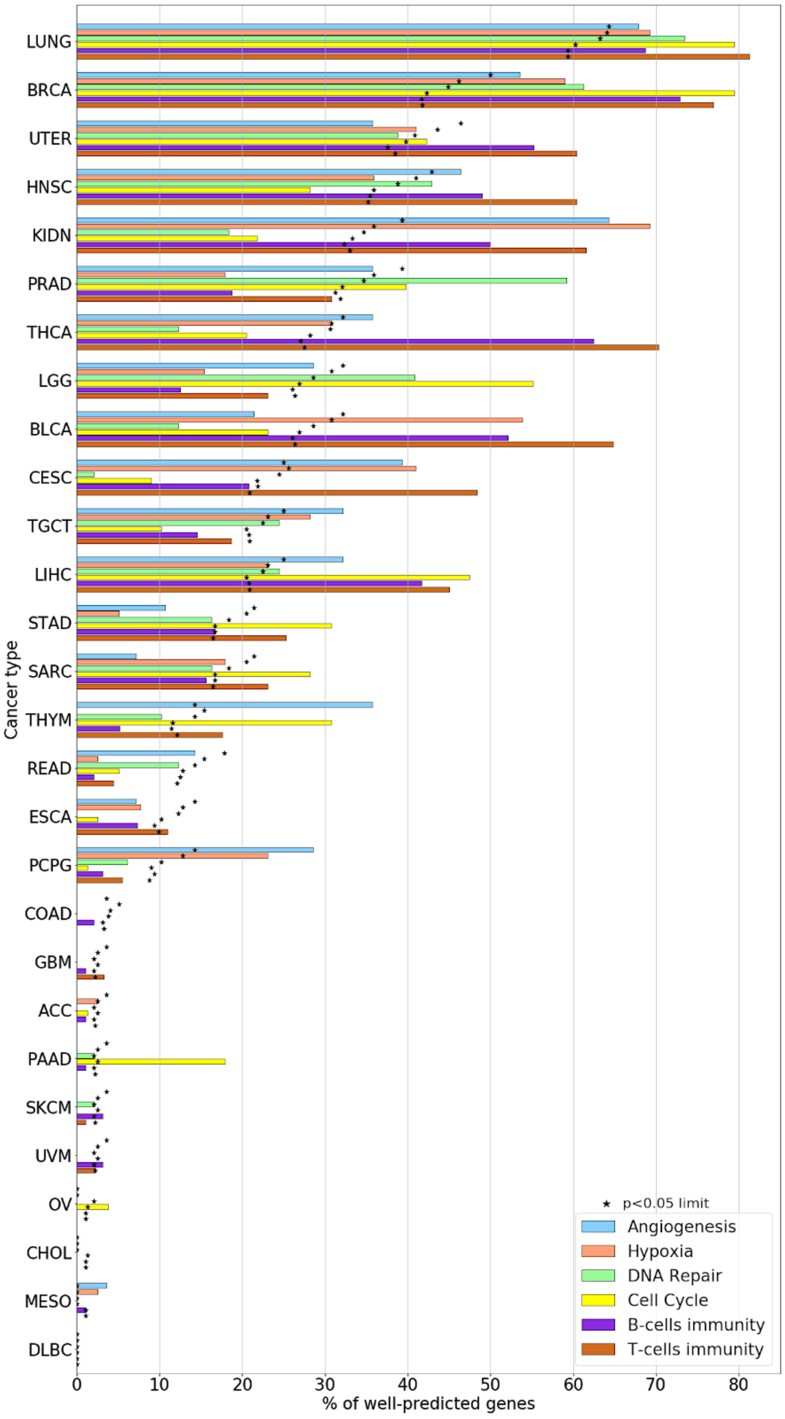

**Figure S3. Percentage of genes for which expression was well-predicted in six common pathways of carcinogenesis.** We show the percentage of genes for which expression was well-predicted (as described in the main text and in Fig.2) for the six studied hallmark pathways for cancer. The black stars indicate the percentage required for each cancer dataset and each pathway to be considered significantly better predicted than a corresponding random list of genes of the same length as the pathway gene list (as explained in the main text and in the Methods section).

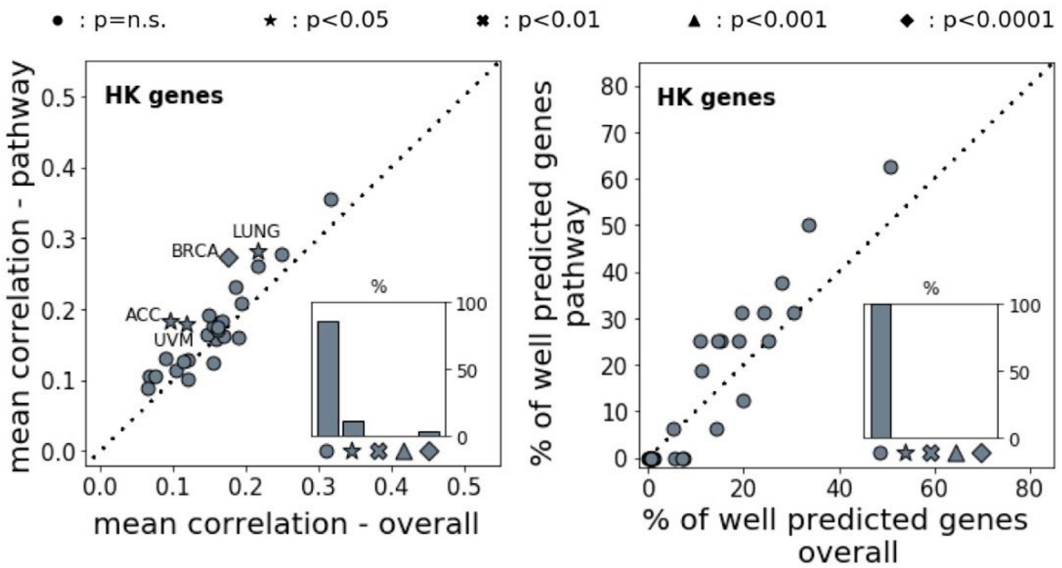

**Figure S4. Prediction for a housekeeping genes signature.** As in Fig.3, for the housekeeping gene signature described in Table S4. **Left panel.** The indicated statistical significance refers to the probability of obtaining a correlation $R > R_p$ in the distribution of correlations for random lists, for each given cancer type. Insets show the percentages of the different cases of statistical significance between cancer types. **Right panel.** As in the left panel, but for the percentage of genes for which expression was of well-predicted (as defined in the text and in Fig. 2). HK = housekeeping.

| Ingenuity Canonical Pathways | -log(p-value) | Ratio | Genes |
|---|---|---|---|
| Th1 and Th2 Activation Pathway | 14.1 | 0.0909 | *CD247, CCR1, IL2RG, CD3E, IL12RB1, HAVCR2, CXCR3, CD8A, CD3D, CD3G, PIK3CG, IL10RA, CD86, IL2RA, VAV1, HLA-DPB1, HLA-DPA1* |
| iCOS-iCOSL Signaling in T Helper Cells | 12.9 | 0.112 | *PTPRC, CD247, CD3G, IL2RG, LCK, CD3E, PIK3CG, ZAP70,TRAT1, CD86, IL2RA, VAV1, CD3D, ITK* |
| Th1 Pathway | 12.4 | 0.102 | *CD247, CD3E, IL12RB1, HAVCR2, CXCR3, CD8A, CD3D, CD3G, PIK3CG, IL10RA, CD86, VAV1, HLA-DPB1, HLA-DPA1* |
| T Cell Receptor Signaling | 10.6 | 0.1 | *PTPRC, CD247, CD3G, LCK, PTPN7, CD3E, PIK3CG, ZAP70, VAV1, CD8A, CD3D, ITK* |
| Th2 Pathway | 10.5 | 0.0855 | *CCR1, CD247, CD3G, IL2RG, CD3E, IL12RB1, PIK3CG, CD86, IL2RA, VAV1, HLA-DPB1, CD3D, HLA-DPA1* |
| CD28 Signaling in T Helper Cells | 10 | 0.0896 | *PTPRC, CD247, CD3G, LCK, CD3E, WAS, PIK3CG, ZAP70, CD86, VAV1, CD3D, ITK* |
| Primary Immunodeficiency Signaling | 8.87 | 0.16 | *PTPRC, IL2RG, LCK, CD3E, ZAP70, CIITA, CD8A, CD3D* |
| CTLA4 Signaling in Cytotoxic T Lymphocytes | 8.84 | 0.099 | *CD247, CD3G, LCK, CD3E, PIK3CG, ZAP70, TRAT1,CD86 CD8A, CD3D* |
| Pathogenesis of Multiple Sclerosis | 8.82 | 0.556 | *CCR1, CCL4, CXCR3, CCL5, CXCL9* |
| Natural Killer Cell Signaling | 7.68 | 0.0752 | *CD247, LCK, SH2D1A, LAIR1, TYROBP, PIK3CG, ZAP70, VAV1, HCST, FCGR3A/FCGR3B* |

**Table S1.** Canonical pathways with the best overlap with the 156 genes for which expression was well-predicted in at least 12 cancer types.

| Ingenuity Canonical Pathways | -log(p-value) | Ratio | Genes |
|---|---|---|---|
| Cell Cycle Control of Chromosomal Replication | 10.6 | 0.286 | *MCM6, CDC45, CDT1, CDK16, CDC6, ORC6, CDC7, CDK1, MCM4, MCM3, MCM2, TOP2A, PRIM2, DBF4, ORC1, MCM7* |
| Mitotic Roles of Polo-Like Kinase | 8.41 | 0.227 | *KIF23, CDC20, PTTG1,PRC1, CDC7, CCNB2, PLK1, CDK1, CCNB1, PLK4, TGFB1, FBXO5, PKMYT1, KIF11, CDC25A* |
| Hepatic Fibrosis / Hepatic Stellate Cell Activation | 8.39 | 0.134 | *COL8A2, CCR5, COL10A1, COL4A2, COL1A2, COL5A1, COL16A1, TIMP1, TGFB1, PDGFRA, TIMP2, CXCL8, COL6A2, FGFR2, MMP2, COL1A1, COL6A3, TGFB3, TGFA, IL10RA, EDNRA, COL11A1, COL9A2, MMP9, COL3A1* |
| Th1 and Th2 Activation Pathway | 7.68 | 0.128 | *CCR5, IL2RG, HLA-DOA, CD3E, IL12RB1, IKZF1, HAVCR2, PIK3R5, HLA-DQA1, LGALS9, FGFR2, SPI1, CD3G, IL18, TGFB1, HLA-DMB, IL10RA, CD86, IL2RA, VAV1, JAG1, JAK3 ,NOTCH1, HLA-DPA1* |
| GP6 Signaling Pathway | 6.85 | 0.141 | *COL8A2,COL6A2,PIK3R5,COL10A1,FGFR2,COL4A2,LAMC2,COL16A1,COL5A1,COL1A2,COL1A1,COL6A3,SYK,LAMB1,FCER1G,COL11A1,COL9A2,LCP2,COL3A1* |
| Th2 Pathway | 6.69 | 0.132 | *CCR5, IL2RG, HLA-DOA, CD3E, IL12RB1, IKZF1, PIK3R5, HLA-DQA1, FGFR2, SPI1, CD3G, TGFB1, HLA-DMB, CD86, IL2RA, VAV1, JAG1, JAK3, NOTCH1, HLA-DPA1* |
| Th1 Pathway | 6.07 | 0.131 | *CCR5, HLA-DOA,CD3E, IL12RB1, HAVCR2, PIK3R5, HLA-DQA1, LGALS9, FGFR2, CD3G, IL18, HLA-DMB, IL10RA, CD86, VAV1, JAK3, NOTCH1, HLA-DPA1* |
| Role of BRCA1 in DNA Damage Response | 5.59 | 0.162 | *RAD51, FANCB, FANCD2, RFC4, FANCG, BARD1, SMARCD1, PLK1, E2F3, BLM, RBL1, E2F2, CHEK1* |
| CD28 Signaling in T Helper Cells | 5.56 | 0.127 | *HLA-DOA, ARPC1B, CD3E, PIK3R5, HLA-DQA1, FGFR2, IKBKE, CD3G, LCK, CARD11, SYK, ITPR3, HLA-DMB, FCER1G, CD86, VAV1, LCP2* |
| iCOS-iCOSL Signaling in T Helper Cells | 5.32 | 0.128 | *HLA-DOA, IL2RG, CD3E, HLA-DQA1, PIK3R5, FGFR2, IKBKE, CD3G, LCK, HLA-DMB, ITPR3, FCER1G, CD86, VAV , IL2RA, LCP2* |

**Table S2.** Canonical pathways with the best overlap with the genes for which expression was best predicted (correlation coefficient above 0.4) in liver hepatocellular carcinomas.

| Ingenuity Canonical Pathways | -log(p-value) | Ratio | Genes |
|---|---|---|---|
| Primary Immunodeficiency Signaling | 16.3 | 0.4 | *CD19, IL2RG, CD3E, IGLL1/IGLL5, CIITA,CD79A, IGHG1, CD8A,TNFRSF13C, CD3D, TAP1, IL7R, LCK, IGHG3, ICOS, ZAP70, IGHM, IGHA1, JAK3, TAP2* |
| Cell Cycle Control of Chromosomal Replication | 12.8 | 0.321 | *MCM5, MCM6, CDC45, CDT1, CDC6, CDC7, ORC6, CDK1, MCM4, MCM3, PCNA, MCM2, TOP2A, PRIM2, CHEK2, DBF4, MCM7, ORC1* |
| Mitotic Roles of Polo-Like Kinase | 10.4 | 0.258 | *KIF23,CDC25C, ESPL1, CDC20, PTTG1, PRC1, CDC7, CCNB2, PLK1, CDK1, CCNB1, CDC25B, PLK4, FBXO5, CHEK2, KIF11, CDC25A* |
| Th1 and Th2 Activation Pathway | 9.73 | 0.144 | *CD247,CD3E,KLRD1,IL12RB1,CXCR3,CD8A,TBX21,IL18R1,IL2RB,RUNX3,IFNG,IL2RG,IKZF1,IL12RB2,CD3D,STAT4,CD3G,LTA,ICOS,GFI1,CXCR6,S1PR1,APH1B,HLA-DOB,IL2RA,PIK3CD,JAK3* |
| Cell Cycle: G2/M DNA Damage Checkpoint Regulation | 9.32 | 0.286 | *CDC25C, CKS2, YWHAZ, CCNB2, PLK1, AURKA, CDK1, CHEK1, SKP2, CCNB1, CDC25B, CKS1B, TOP2A, CHEK2* |
| Role of CHK Proteins in Cell Cycle Checkpoint Control | 8.37 | 0.246 | *CDC25C, PLK1, E2F3, CDK1, CHEK1, PCNA, RFC4, E2F1, RFC2, CLSPN, E2F2, CHEK2, E2F8, CDC25A* |
| Estrogen-mediated S-phase Entry | 8.24 | 0.385 | *CCNA2, CCNE1, E2F1, E2F3, ESR1, E2F8, E2F2, CDK1, SKP2, CDC25A* |
| Th2 Pathway | 8.07 | 0.145 | *CD247, RUNX3, IFNG, IL2RG, CD3E, IL12RB1, IKZF1, IL12RB2, TBX21, CD3D, STAT4, CD3G, ICOS, GFI1, CXCR6, S1PR1, APH1B, HLA-DOB, IL2RA, PIK3CD, JAK3, IL2RB* |
| Th1 Pathway | 7.46 | 0.146 | *CD247, RUNX3, IFNG, CD3E, IL12RB1, KLRD1, CXCR3, IL12RB2, TBX21, CD8A, CD3D, IL18R1, STAT4, CD3G, LTA, ICOS, APH1B, HLA-DOB, PIK3CD, JAK3* |
| T Helper Cell Differentiation | 6.9 | 0.192 | *IL6ST, IFNG, IL2RG, IL12RB1, IL21R, FOXP3, IL12RB2, TBX21, IL18R1, STAT4, ICOS, HLA-DOB, IL2RA,TNFRSF1B* |

**Table S3.** Canonical pathways with the best overlap with the genes for which expression was best predicted (correlation coefficient above 0.4) in breast cancer samples.

| Angio genesis | Hypoxia | DNA repair | Cell cycle | B cells | T cells | Housekee ping |
|---|---|---|---|---|---|---|
| *CDC42* | *ADM* | *CETN2* | *ATM* | *MASP2* | *CD79B* | *RPL32* |
| *FGFR1* | *ALDOA* | *DDB1* | *ATR* | *IGLC7* | *BTLA* | *PPIA* |
| *FLT1* | *BHLHE40* | *DDB2* | *BUB1* | *IGLC3* | *WAS* | *PGK1* |
| *FLT4* | *BNIP3L* | *ERCC1* | *BUB1B* | *IGKV3-20* | *CTSC* | *HMBS* |
| *HIF1A* | *CA12* | *ERCC2* | *BUB3* | *C3* | *ZAP70* | *GAPDH* |
| *HRAS* | *CCNG2* | *ERCC3* | *CCNA1* | *IGHG4* | *FYN* | *GUSB* |
| *ITGAV* | *CDKN1A* | *ERCC4* | *CCNA2* | *IGKV4-1* | *ANXA1* | *TBP* |
| *JAG1* | *CDKN1B* | *ERCC5* | *CCNB1* | *CD74* | *IFNG* | *PSMB2* |
| *KDR* | *COL5A1* | *ERCC8* | *CCNB2* | *C8A* | *PVR* | *ALB* |
| *MAPK14* | *CP* | *FEN1* | *CCND1* | *IGLL1* | *C3* | *HPRT1* |
| *NRP1* | *DDIT3* | *GTF2H1* | *CCND2* | *MLH1* | *LILRB1* | *EMC7* |
| *NOS3* | *EDN2* | *GTF2H3* | *CCND3* | *SERPING1* | *HRAS* | *RPS27* |
| *NFATC4* | *ENO1* | *GTF2H5* | *CCNE1* | *POU2F2* | *CD74* | *RPLP0* |
| *PIK3CA* | *F3* | *LIG1* | *CCNE2* | *LIG4* | *TRAF6* | *SDHA* |
| *PIK3CB* | *FOS* | *MPG* | *CCNH* | *HLA-DRB1* | *LILRB5* | *ACTB* |
| *PIK3CG* | *GAPDH* | *PCNA* | *CDC14A* | *CFI* | *HLA-B* | *AC010970.1* |
| *PIK3R1* | *HIF1A* | *POLB* | *CDC16* | *IL4R* | *FCGR1B* | |
| *PDGFA* | *IGFBP3* | *POLD1* | *CDC20* | *CD40LG* | *PRKCQ* | |
| *PRKCA* | *HK1* | *POLD3* | *CDC23* | *BCL3* | *HLA-DRB1* | |
| *PRKCB* | *HK2* | *POLD4* | *CDC25A* | *IGLC6* | *ERAP1* | |
| *PLCG1* | *HMOX1* | *POLH* | *CDC25B* | *RNF8* | *IL4R* | |
| *PTK2* | *IGFBP1* | *POLL* | *CDC25C* | *IGHM* | *RIPK2* | |
| *PTGS2* | *IL6* | *POLR2A* | *CDC26* | *HLA-DQB1* | *IL4* | |
| *PXN* | *JUN* | *POLR2C* | *CDC27* | *C4BPB* | *CD40LG* | |
| *SHC1* | *LDHA* | *POLR2D* | *CDC45* | *BATF* | *HMGB1* | |

| | | | | | |
|---|---|---|---|---|---|
| *SH2D2A* | *MIF* | *POLR2E* | *CDC6* | *ZP3* | *KLRK1* |
| *VEGFA* | *P4HA1* | *POLR2F* | *CDC7* | *IGHG3* | *CTLA4* |
| *VAV2* | *PDGFB* | *POLR2G* | *CDK1* | *SWAP70* | *LILRB2* |
| | *PFKL* | *POLR2H* | *CDK2* | *C1QBP* | *PTPRC* |
| | *PFKP* | *POLR2I* | *CDK4* | *BCL10* | *HAVCR2* |
| | *PGF* | *POLR2J* | *CDK6* | *FAS* | *C4BPB* |
| | *PGK1* | *POLR2K* | *CDK7* | *EXO1* | *SLAMF1* |
| | *PLAUR* | *RAD51* | *CDKN1A* | *IGLC2* | *ZP3* |
| | *SLC2A1* | *RAD52* | *CDKN1B* | *IGHG2* | *LILRA1* |
| | *SLC2A3* | *RBX1* | *CDKN2A* | *IGKV2-40* | *TRAT1* |
| | *STC1* | *REV3L* | *CDKN2B* | *CD55* | *STAT6* |
| | *TGFB3* | *RFC2* | *CDKN2C* | *PRKCD* | *PTPN6* |
| | *TGM2* | *RFC3* | *CDKN2D* | *INPP5D* | *CTSS* |
| | *VEGFA* | *RFC4* | *CHEK1* | *AICDA* | *BCL10* |
| | | *RFC5* | *CHEK2* | *IGHV4OR15-8* | *LYN* |
| | | *RPA2* | *CUL1* | *IL13RA2* | *TGFB1* |
| | | *RPA3* | *DBF4* | *C4A* | *TAP2* |
| | | *TP53* | *E2F1* | *C1R* | *TRPM4* |
| | | *XPC* | *E2F2* | *IGHA2* | *CD8A* |
| | | *ATM* | *E2F3* | *C4BPA* | *SYK* |
| | | *ATR* | *E2F4* | *NBN* | *CTSH* |
| | | *CHEK1* | *E2F5* | *IGHG1* | *PIK3CD* |
| | | *CHEK2* | *HDAC1* | *C7* | *FOXP3* |
| | | *H2AFX* | *KI67* | *MBL2* | *CTSL* |
| | | | *MAD1L1* | *IGHV2-70* | *PAG1* |
| | | | *MAD2L1* | *C1QA* | *NECTIN2* |
| | | | *MCM2* | *HSPD1* | *CD79A* |

| | | | | | |
|---|---|---|---|---|---|
| | | | *MCM3* | *C1RL* | *C4BPA* |
| | | | *MCM4* | *IGKV1D-33* | *TNFSF18* |
| | | | *MCM5* | *C9* | *CRTAM* |
| | | | *MCM6* | *MSH6* | *BTK* |
| | | | *MCM7* | *C8G* | *IL2* |
| | | | *MDM2* | *IGLL5* | *RAET1E* |
| | | | *MYC* | *IGLV7-43* | *HSPD1* |
| | | | *ORC1* | *IRF7* | *HLA-A* |
| | | | *PCNA* | *CD27* | *MSH6* |
| | | | *PKMYT1* | *GCNT3* | *TNFRSF13 C* |
| | | | *PLK1* | *C6* | *EIF2AK4* |
| | | | *PTTG1* | *ERCC1* | *HLA-E* |
| | | | *RAD21* | *IGKV3D-20* | *IRF7* |
| | | | *RB1* | *EXOSC3* | *PRKCB* |
| | | | *RBL1* | *FCER1G* | *CD4* |
| | | | *RBL2* | *C1S* | *EXOSC3* |
| | | | *SKP1* | *IGKV1-5* | *IL12B* |
| | | | *SKP2* | *MSH2* | *FCER1G* |
| | | | *SMC1A* | *IGKC* | *JAK3* |
| | | | *SMC1B* | *GAPT* | *B2M* |
| | | | *SMC3* | *C1QC* | *BCL6* |
| | | | *STAG1* | *CR2* | *LILRB3* |
| | | | *STAG2* | *RNF168* | *MEF2C* |
| | | | *TFDP1* | *IGHA1* | *MALT1* |
| | | | *TP53* | *IGHV3-23* | *LILRB4* |
| | | | *WEE1* | *HLA-DRB5* | *CSK* |
| | | | | *CR1* | *TAP1* |

| | | | | | |
|---|---|---|---|---|---|
| | | | | *IGKV3D-11* | *HLA-DRB5* |
| | | | | *TRDC* | *CR1* |
| | | | | *TLR8* | *GATA3* |
| | | | | *EXOSC6* | *CD8B* |
| | | | | *C5* | *CD86* |
| | | | | *C2* | *SUSD4* |
| | | | | *SLA2* | *MAP3K7* |
| | | | | *SUSD4* | *SLC11A1* |
| | | | | *CD46* | *NLRP10* |
| | | | | *CLU* | *IFNB1* |
| | | | | *C8B* | *ORAI1* |
| | | | | *C4B* | *ITK* |
| | | | | *IGHV1OR21-1* | |
| | | | | *IGHE* | |
| | | | | *C1QB* | |
| | | | | *IGHD* | |
| | | | | *IGLC1* | |

**Table S4.** List of genes for the six signatures chosen for angiogenesis, hypoxia, DNA repair, cell cycle pathways, B-cell and T-cell immune responses.
